# Cross-Protective Antibodies Against Common Endemic Respiratory Viruses

**DOI:** 10.1101/2022.06.26.497647

**Authors:** Madelyn Cabán, Justas V. Rodarte, Madeleine Bibby, Matthew D. Gray, Justin J. Taylor, Marie Pancera, Jim Boonyaratanakornkit

## Abstract

Respiratory syncytial virus (RSV), human metapneumovirus (HMPV), and human parainfluenza virus types one (HPIV1) and three (HPIV3) are a major cause of death, morbidity, and health care costs worldwide, and they can exact a significant toll on immunocompromised patients, the elderly, and those with underlying lung disease. There is an unmet medical need for safe and effective medications for many of the viruses responsible for common respiratory viral infections in vulnerable patient populations. While a protective monoclonal antibody exists for RSV, clinical use is limited to high-risk infant populations. Here, we present the discovery, *in vitro* characterization, and *in vivo* efficacy testing of two cross-neutralizing monoclonal antibodies, one targeting both HPIV3 and HPIV1 and the other targeting both RSV and HMPV. The 3×1 antibody is capable of targeting multiple parainfluenza viruses; the MxR antibody shares features with other previously reported monoclonal antibodies that are capable of neutralizing both RSV and HMPV. We obtained structures using cryo-electron microscopy of these antibodies in complex with their antigens to 3.62 Å resolution for 3×1:HPIV3 and to 2.24 Å for MxR:RSV, providing a structural basis to corroborate our *in vitro* characterization of binding and neutralization. Together, a cocktail of 3×1 and MxR could have clinical utility in providing broad protection against four of the respiratory viruses that cause significant morbidity and mortality in at-risk individuals.

## INTRODUCTION

Respiratory viruses are a major cause of death worldwide, with an estimated 2.7 million attributable deaths in 2015^1^. While a vaccine to prevent RSV infection may be on the horizon^2, 3^, protective vaccines for HMPV, HPIV3, and HPIV1 have not been developed. Even if protective vaccines existed for these four respiratory viruses, vaccination of highly immunocompromised individuals rarely achieves protective immunity. Additionally, vaccination prior to immune-ablative therapies is often ineffective or wanes quickly, failing to maintain durable protection^4, 5, 6^. Together, RSV, HMPV, HPIV1, and HPIV3 represent a serious threat to immunocompromised patients and, prior to the COVID-19 pandemic, were responsible for the majority of viral lower respiratory infections in hematopoietic stem cell transplant recipients^7, 8^. In adults with other risk factors, the burden of disease from HMPV and the parainfluenza viruses is also comparable to RSV^9, 10^. Further, with the exception of rhinoviruses, RSV, HMPV, and the parainfluenza viruses also collectively account for most of the respiratory viruses identified in hospitalized adults prior to 2020^11, 12^.

Although mitigation strategies during the COVID-19 pandemic such as masking, social distancing, and shut-downs led to declines in cases of other respiratory viruses during the 2020-2021 cold and flu seasons, cases of RSV, HMPV, and HPIVs are beginning to surge again and are expected to return to pre-pandemic levels of circulation in the next few years^9, 10^. In fact, models project large future outbreaks of non-SARS-CoV-2 respiratory viruses due to an increase in size of the susceptible population following a period of reduced spread^13^. Additionally, since endemic respiratory viruses tend to circulate seasonally, co-infections with more than one respiratory virus can occur and have been associated with worse outcomes in vulnerable populations^14, 15, 16^.

The administration of neutralizing monoclonal antibodies (mAbs) provides an effective alternative to vaccination to protect against viral infections. Although the anti-RSV mAb palivizumab received FDA-approval in 1998 as prophylaxis in high-risk infants^17^, it remains relatively unused in older immunocompromised children or adults. Since the approval of palivizumab, even more potent mAbs against RSV have progressed through clinical trials, with the primary goal of replacing palivizumab as the standard of care for prophylaxis in high-risk infants. This focus has been driven in part because RSV causes up to 80% of bronchiolitis in infants^18, 19^. However, in immunocompromised adults, the respiratory virus landscape is much more heterogeneous ^7, 8^. Therefore, an effective strategy to reduce the overall burden of the broad range of lower respiratory tract infections in at-risk adults and immunocompromised patients must rely on targeting multiple viruses simultaneously, rather than a single virus. Despite advances in research on RSV prevention, the role of passive immunization for other respiratory viruses remains poorly defined and no mAbs are currently available in the clinic that can prevent HMPV, HPIV1, or HPIV3 infection.

To efficiently achieve broader protection against these viruses, we sought to identify cross-neutralizing mAbs that could target more than one virus at a time. RSV, HMPV, HPIV3, and HPIV1 all produce class I fusion (F) proteins which are essential surface glycoproteins specialized to mediate fusion between viral and host cell membranes during viral entry. HPIV1 and HPIV3 belong to the same *Respirovirus* genus, and their F sequences share 65% amino acid sequence homology. RSV and HMPV belong to the same *Pneumoviridae* family, and their F sequences share approximately 54% homology. The F proteins transition between a metastable prefusion (preF) conformation and a stable postfusion (postF) conformation^20, 21^. Since preF is the major conformation on infectious virions, antibodies to preF tend to be the most potent at neutralizing virus^22, 23, 24, 25^. In the present study, we leveraged the homology between the related F proteins and their preF conformations to identify and clone two potent cross-neutralizing mAbs, 3×1 and MxR, from human memory B cells. 3×1 cross-neutralizes both HPIV3 and HPIV1, while MxR effectively cross-neutralizes both HMPV and RSV. Together, 3×1 and MxR comprise an antibody cocktail with the ability to achieve simultaneous protection against multiple viruses which could be beneficial to at-risk populations who are at a significant immunological disadvantage when infected with respiratory viruses.

## RESULTS

### Identification of a novel cross-neutralizing antibody against multiple parainfluenza viruses

Since virtually all humans have been exposed to RSV, HMPV, HPIV3 and HPIV1^26^, it was not necessary to pre-screen donors for seropositivity. Because there is a lack of mAbs currently under development against the parainfluenza viruses, we first focused our efforts on screening for HPIV3/HPIV1 cross-neutralizing B cells. Since we were unable to produce the HPIV1 F protein in the preF conformation, to conduct this screen we used the F protein of HPIV3 alone by leveraging a “bait-and-switch” strategy (**Fig. 1a, b**). Here, HPIV3-binding B cells were isolated by incubating 200 million human splenocytes with tetramers of HPIV3 preF conjugated to APC and tetramers of HPIV3 postF conjugated to APC/Dylight755 followed by magnetic enrichment with anti-APC microbeads. Nine-hundred B cells that bound HPIV3 preF tetramers but not postF tetramers were individually sorted into wells containing CD40L/IL2/IL21-producing 3T3 feeder cells and then incubated for 13 days to stimulate antibody secretion into culture supernatants. To identify wells containing candidate B cells expressing cross-neutralizing B cell receptors, culture supernatants from the individually sorted HPIV3-binding B cells were mixed with live HPIV1 virus and screened for their ability to reduce plaque development (**Fig. 1a, c**). Two out of the 900 HPIV3-binding B cells produced antibodies that could neutralize HPIV1 (**Fig. 1c**). From these cells, the heavy (VH3-23) and light (Vλ3-19) chain alleles were sequenced and cloned successfully to produce a mAb with cross-neutralizing capability against HPIV3/HPIV1 which we named 3×1 (**Fig. 1b, c**). 3×1 was isolated from a B cell expressing the IgA isotype. Since palivizumab and most other mAbs being developed against respiratory viruses utilize an IgG1 constant region, 3×1 was also cloned and produced for further study as an IgG1.

**Figure 1.**
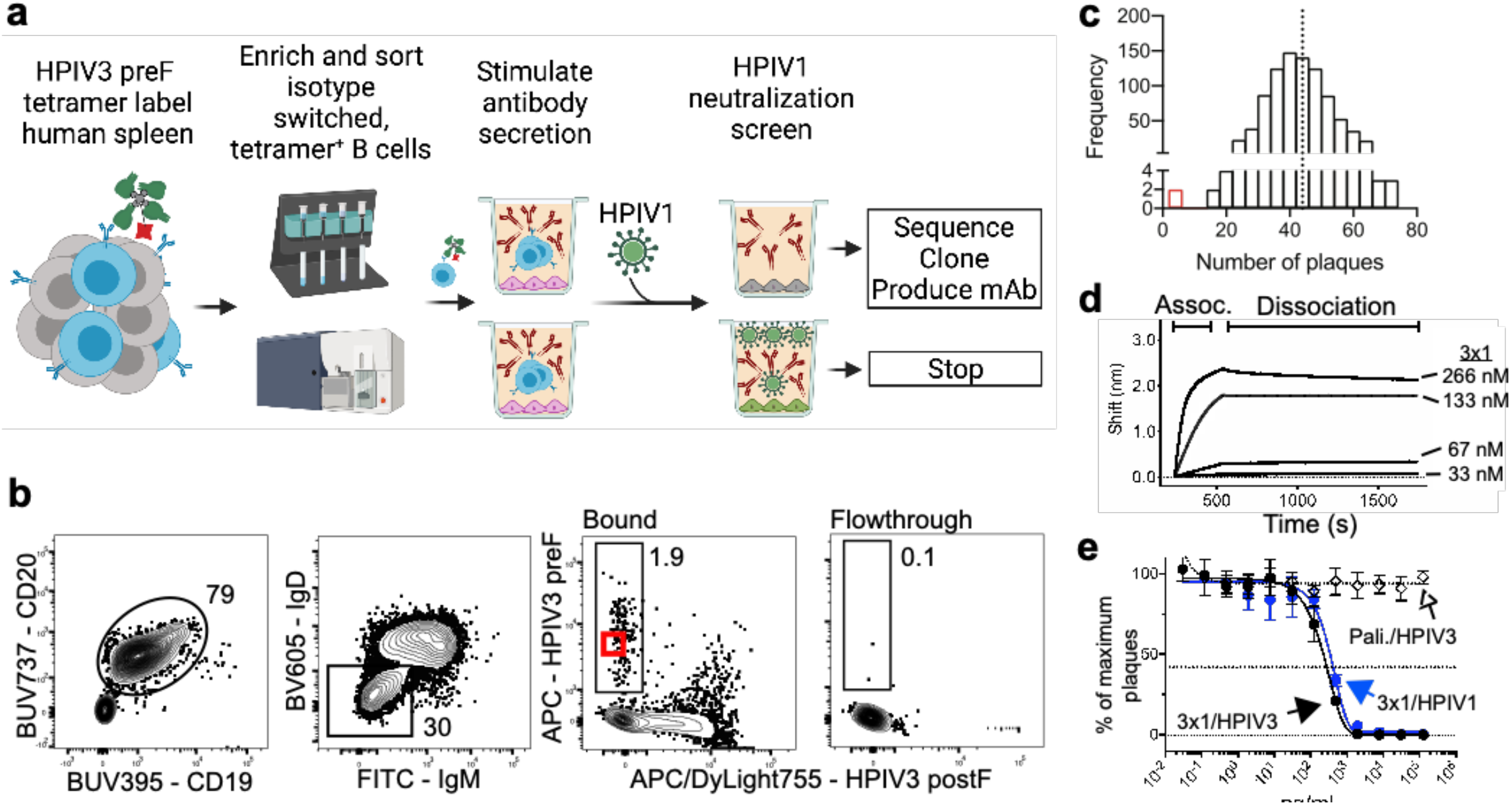
Identification of a novel HPIV3/HPIV1 cross-neutralizing monoclonal antibody. (**a**) “Bait-and-switch” approach using a single antigen to identify B cells that cross-neutralize another related virus. HPIV3-binding B cells from human spleen were labeled with APC-conjugated tetramers of HPIV3 prefusion protein (preF). (**b**) Flow cytometry plot of HPIV3 preF-binding B cells after gating for live, CD3^-^CD14^-^CD16^-^CD19^+^CD20^+^ (B cells), IgD^-^/IgM^-^ (isotype-switched), and APC/Dylight755^-^ HMPV postF^-^ (to exclude cells binding to HPIV3 postF, APC, or streptavidin). The bound fraction contains cells magnetically enriched cells using APC-specific microbeads. Numbers on plots are percentages of total cells in the gate. The red box indicates the B cell from which 3×1 mAb was derived. (**c**) Frequency distribution of HPIV1 plaques per well from the neutralization screen. The dotted line indicates the mean number of plaques in negative control wells containing virus in the absence of antibodies. (**d**) Apparent affinity (*K*_D_) of 3×1 to HPIV3 preF was measured by biolayer interferometry (BLI). Association with 3×1 to HPIV3 preF-loaded probe was measured for 300 s followed by dissociation for 1200 s. Measurements are normalized against an isotype control antibody. (**e**) Vero cells were infected with HPIV3 or HPIV1 in the presence of serial dilutions of palivizumab or 3×1. The dotted midline indicates the PRNT_60_. Data points are the average +/- SD from three independent experiments.

Although we did not have the F protein of HPIV1 stabilized in the preF conformation, we did estimate the apparent binding affinity between 3×1 and the F protein of HPIV3 in the preF conformation. 3×1 bound extremely well to the preF protein of HPIV3 (*K*_D_ < 10^−12^ M) (**Fig. 1d**). The neutralizing potency of 3×1 was determined by a 60% plaque reduction neutralization test (PRNT_60_) using live virus to infect Vero cells. 3×1 had a similarly high neutralization potency against both HPIV1 and HPIV3, with a PRNT60 of 352 and 242 ng/mL, respectively (**Fig. 1e**). These results suggest that the epitope of 3×1 might be functionally conserved between HPIV3 and HPIV1.

### Cryo-EM structure of 3×1 Fab in complex with HPIV3 preF

Since the antigenic landscape of HPIV3 preF is not well characterized, we first performed cross-competition binding experiments to gauge the antigenic sites on HPIV3 preF allowing for neutralization. The epitope for the cross-neutralizing mAb 3×1 did not appear to overlap with the epitopes of mAbs that we had previously isolated which bind to the apex of preF and neutralized HPIV3 (**Fig. 2a**)^27^. However, the epitope of 3×1 did appear to overlap with the epitope of a previously described antibody PI3-A12, which binds to a novel antigenic site we named site X^27^. To determine how 3×1 interacts with site X, we obtained a cryo-EM structure of one 3×1 Fab in complex with HPIV3 preF resolved to 3.62 Å (**Fig. S1a and Table S1**). We also obtained a structure showing three Fabs bound to HPIV3; this map was limited to 4.3 Å resolution, and consequently was not used for model building (**Fig. S2a**). 3×1 binds each vertex within domain III of HPIV3 preF, protruding perpendicularly and binding only one protomer, with no additional contacts to other regions (**Fig. 2b**). Only four of the six CDRs are involved in binding, with CDRH1 and CRDL2 being too short to contact the glycoprotein surface (**Fig. 2c, f**). 3×1 binds at site X with a total buried surface area (BSA) of ∼812 Å^2^, of which the VH contributes ∼612 Å^2^ and the VL contributes ∼200 Å^2^ (**Fig. 2b and Fig. S3**). 3×1 neutralizes both HPIV3 and HPIV1, whereas PI3-A12 neutralizes only HPIV3^27^. 3×1 shares little CDR sequence similarity with PI3-A12 (**Fig. S3c**). When compared to PI3-A12, 3×1 binds the same site but is rotated (**Fig. S3a, b**). Due to the lack of a high-resolution structure for PI3-A12, we could not determine if the 3×1 LC overlaps with the PI3-A12 LC or HC. 3×1 and PI3-A12 may therefore bind distinct epitopes within the same site of HPIV3 preF, as they display significant sequence variation at key residues in 3×1 that facilitate binding (**Fig. S3c**).

**Figure 2.**
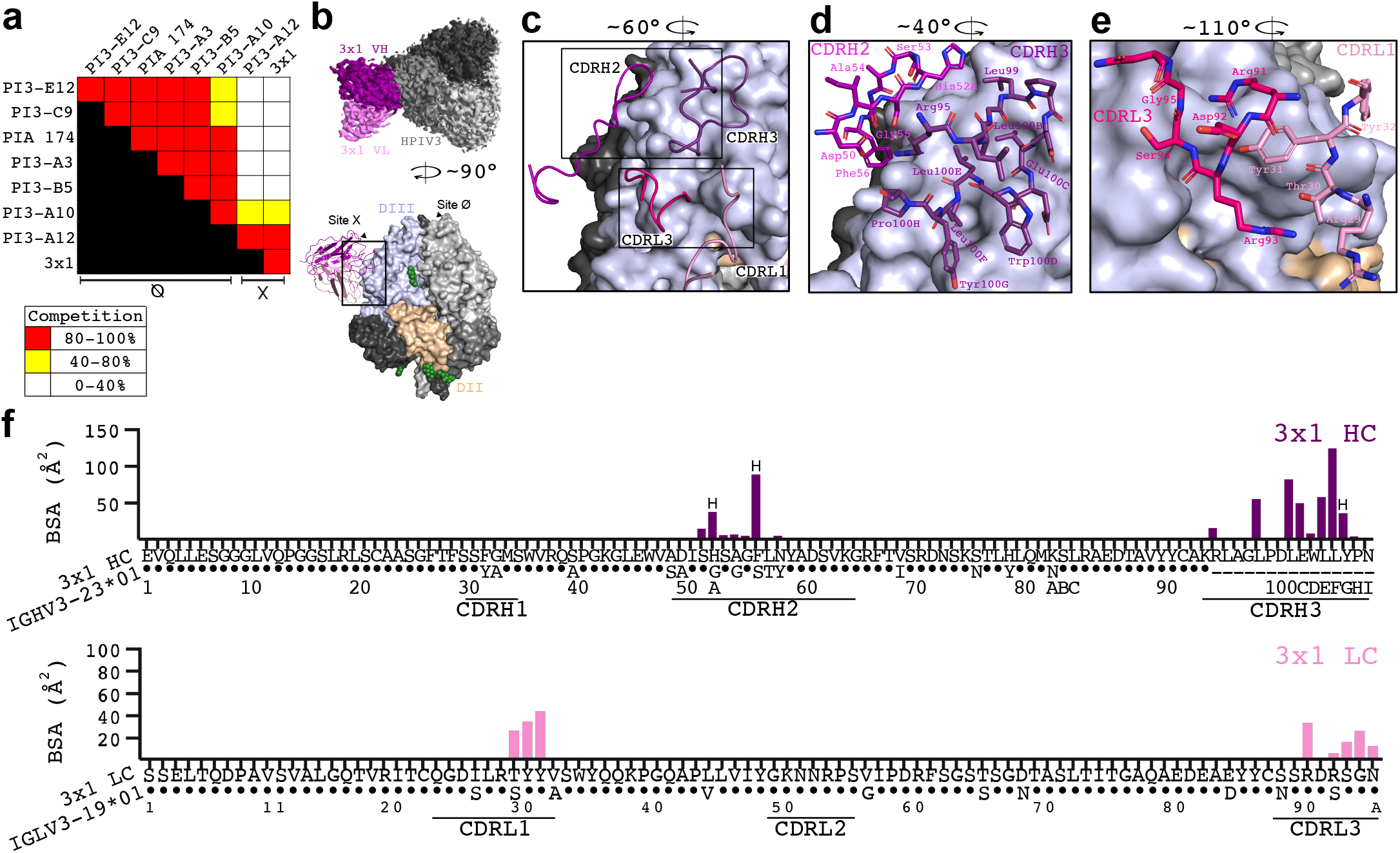
Analysis of the HPIV3 preF epitope bound by 3×1. (**a**) BLI measurement of the ability of the mAb listed on the left side of the chart to prevent binding of the mAb listed on the top. Expressed as the percent drop in maximum signal compared to the maximum signal in the absence of competing mAb. (**b**) Cryo-EM structure of 3×1 Fab in complex with HPIV3 preF. Top, a top-down view of the 3×1:HPIV3 map is shown with one Fab, with the 3×1 V_H_ shown in purple, 3×1 V_L_ in pink, and HPIV3 preF in shades of grey. Bottom, a surface representation of the complex is shown rotated 90 degrees. Glycans are shown as green spheres. (**c**) Detailed view of the 3×1 binding site with only the interacting CDRs shown in cartoon, rotated 60 degrees from panel (**b**). Boxes show locations of panels (**d**) and (**e**). (**d**) Zoomed in view of the CDRH3 binding site, rotated 40 degrees from panel (**b**). (**c, d**) CDR residues which make no contacts with the spike have been hidden from view for clarity. (**e**) Zoomed in view of the CDRL3 binding site, rotated 100 degrees from panel (**b**). (**f**) BSA plots for each residue that interacts with the preF protomer, atop a sequence alignment to the germline V_H_ and V_L_ for 3×1.

Further analysis of the local resolution of our map indicates the binding site had a higher resolution of ∼3.0 Å compared to the overall resolution of 3.62 Å (**Fig. S1a**). The 3×1 VH and VL together specifically bind the cleft between the HRA helix and a sheet-turn-sheet motif, a short contiguous region of HPIV3 (**Fig. S2b-d and S4**). This region is crucial for the preF to postF rearrangement, with both the HRA helix and sheet-turn-sheet motif displaying > 9 Å of movement during rearrangement^28^. Due to the significant motion required of this site during fusion, 3×1 binding at this location presents a strong structural basis for the high neutralizing potency of 3×1 against HPIV3. The CDRH2 and CDRH3 of 3×1 form several protrusions into grooves on the surface of HPIV3 preF using non-polar residues (**Fig. 2c, d**). His52A_HC_ and Phe56_HC_ (CDRH2) along with Leu100B_HC_ and Leu100F_HC_ (CDRH3) are four residues which account for ∼55 % of the total VH BSA (**Fig. 2f**). Most other contacting residues within the VH are polar residues which form contacts with generally < 50 Å^2^ of BSA. The VL residues of 3×1 form contacts with HPIV3 preF using polar functional groups and the CDR loop backbone, with CDRL1 encompassing a large protrusion on HPIV3 preF (**Fig. 2e, f**). This CDRL1 extension is supported by Tyr31_LC_ and Arg91_LC_ which form an intra-Fab hydrogen bond, with Arg91_LC_ forming an additional bond to HPIV3 preF at Asp143, which is conserved in HPIV1 (**Fig. S2e and S5a**).

### Identification of a potent cross-neutralizing antibody against RSV and HMPV

Since RSV and HMPV also contribute significantly to disease in vulnerable patients, we next sought to identify potential HMPV/RSV cross-neutralizing B cells, with the goal of combining a cross-neutralizing mAb against HMPV/RSV with 3×1 in order to create a potent cocktail with expanded breadth. Although many mAbs targeting RSV and HMPV are already in development, we used our approach leveraging fluorescent tetrameric probes to identify unique B cells able to bind recombinant F proteins from RSV and HMPV in the preF, but not in the postF, conformation. RSV preF conjugated to allophyocyanin (APC), HMPV preF conjugated to phycoerythrin (PE), RSV postF conjugated to APC/DyLight755, and HMPV postF conjugated to PE/Dylight650 were mixed with 200 million peripheral blood mononuclear cells (PBMCs) prior to magnetic enrichment with anti-PE and anti-APC microbeads. We then performed fluorescent activated cell sorting on the enriched fraction to isolate isotype-switched B cells binding to the F proteins of RSV and HMPV in the preF conformation but not in the postF conformation, followed by single cell sequencing and cloning of their B cell receptors (**Fig. 3a**). Using this method, the heavy (VH3-21) and light (Vλ1-40) chain alleles of a B cell capable of binding both RSV and HMPV were sequenced and cloned as a mAb which we named MxR (**Fig. 3b**). This B cell expressed the IgG isotype. Like 3×1, MxR was cloned and produced as an IgG1 for further study.

**Figure 3.**
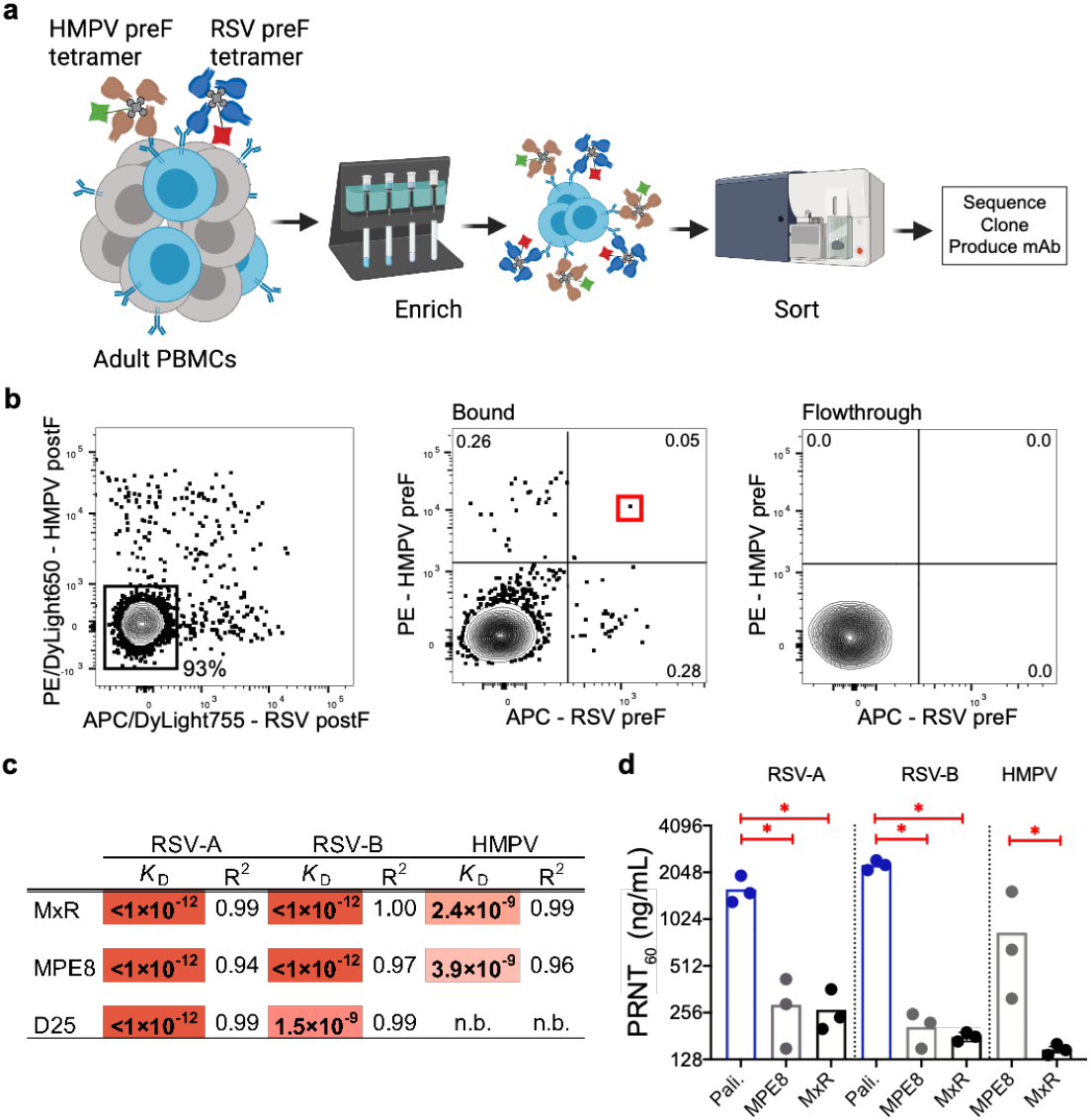
Identification and analysis of HMPV/RSV cross-neutralizing monoclonal antibodies. (**a**) RSV- and HMPV-binding B cells from human blood were labeled with APC-conjugated streptavidin tetramers of biotinylated RSV prefusion protein (preF) and PE-conjugated streptavidin tetramers of biotinylated HMPV preF. (**b**) Flow cytometry plot of RSV and HMPV preF-binding B cells after gating for live, CD3^-^CD14^-^CD16^-^ CD19^+^CD20^+^ (B cells), IgD^-^/IgM^-^ (isotype-switched), and APC/Dylight755^-^ HMPV postF^-^ and PE/DL650-RSV postF^-^ (to exclude cells binding to RSV/HMPV postF, APC, PE, or streptavidin). The bound fraction contains cells magnetically enriched cells using APC-specific microbeads. Numbers on plots are percentages of total cells in the gate. The red box indicates the B cell from which MxR mAb was derived. (**c**) Apparent affinity (*K*_D_) of MxR, MPE8, and D25 measured by BLI. (**d**) Vero cells were infected with RSV-A, RSV-B, or HMPV in the presence of serial dilutions of palivizumab (Pali.), MPE8, or MxR. Data points represent the 60% plaque reduction neutralization titer and are from three independent experiments. The asterisks indicate a *p* < 0.01 using a Mann Whitney two-tailed test.

We compared the apparent binding affinity of MxR to the RSV-specific monoclonal antibody D25, which is being developed under the name nirsevimab for prophylaxis of RSV in infants. RSV has two antigenically distinct subtypes, A and B, which share 91% amino acid sequence homology within the F protein. MxR bound irreversibly with high apparent affinity to the preF proteins of both RSV subtypes A and B (*K*_D_ < 10^−12^ M each), even when the dissociation time was extended to 1,200 seconds (**Fig. 3c and Fig. S6a, d**). The previously reported cross-neutralizing monoclonal antibody MPE8^29^ also exhibited high apparent affinity for both RSV subtypes A and B (**Fig. 3c and Fig. S6b, e**). D25 also bound strongly to the preF protein from RSV subtype B, but with approximately 1,500-fold lower apparent affinity (*K*_D_ = 1.5 × 10^−9^ M) compared to its binding to subtype A (**Fig. 3c and Fig. S6c, f**). MxR also could bind to the preF protein of HMPV (**Fig. 3c and Fig. S6g**), which was expected given the deliberate selection of B cells binding both RSV and HMPV during the sort. Compared to MPE8, the apparent binding affinity of MxR for HMPV is approximately 1.6-fold stronger (**Fig. 3c and Fig. S6g, h**).

Since both subtypes of RSV circulate globally, it was important to assess the neutralization potency (PRNT_60_) of MxR for subtypes A and B. We also compared the neutralizing potency of MxR with the RSV-specific monoclonal antibody palivizumab, which is currently approved for RSV prophylaxis in high-risk infants. We found that MxR neutralized RSV subtype A with at least 6-fold greater potency as compared to palivizumab (**Fig. 3d and Fig. S7a**). In contrast to palivizumab, which has similar potency against both subtypes^30^, MxR was also exceptionally potent against subtype B, with a 12-fold greater potency (**Fig. 3d, Fig. S7b**). MxR and MPE8 had similar potency in neutralizing both RSV subtypes A and B. However, MxR had at least 5-fold greater potency against HMPV, with a PRNT_60_ of 148 ng/mL compared to 838 ng/mL for MPE8 (**Fig. 3d and Fig. S7c**). Based on the concentration of antibody needed to neutralize 60% of live virus, the potency of MxR against HMPV exceeded the relative potency of palivizumab against RSV (**Fig. 3d**).

### Cryo-EM structure of MxR Fab in complex with RSV preF

To better understand the basis for the cross-neutralization observed with MxR, we obtained a cryo-EM structure of three MxR Fabs bound to RSV preF to 2.24 Å resolution (**Fig. S1b and Table S1**). MxR binds primarily to antigenic site III on RSV with an equatorial arrangement of Fabs around RSV preF. Antigenic site III is a quaternary epitope at the junction of domains I and III of one F protomer and domain II of the clockwise adjacent F protomer (referred to as II’) (**Fig. 4a**). MxR binds with a total BSA of ∼1094 Å^2^, with VH contributing ∼694 Å^2^ and the VL contributing ∼400 Å^2^, of which ∼298 Å^2^ contacts with the main F protomer and ∼102 Å^2^ contacts with the adjacent F protomer. This mode of binding is almost identical to the previously reported cross-neutralizing monoclonal antibody MPE8^29^ and the infant monoclonal antibody ADI19425^31^, which are derived from the same germline heavy (IGHV3-21*01) and light chain (IGLV1-40*01) alleles. Comparison of the per-residue BSA of RSV preF in complex with MxR, MPE8, and ADI-19425 revealed common binding regions that share high sequence homology with HMPV (**Fig. S5b**). The sequence and structure of both CDR1s and CDR2s are nearly identical across all three antibodies, with MxR overall having more mutations from germline than the other two (**Fig. 4b, e**).

**Figure 4.**
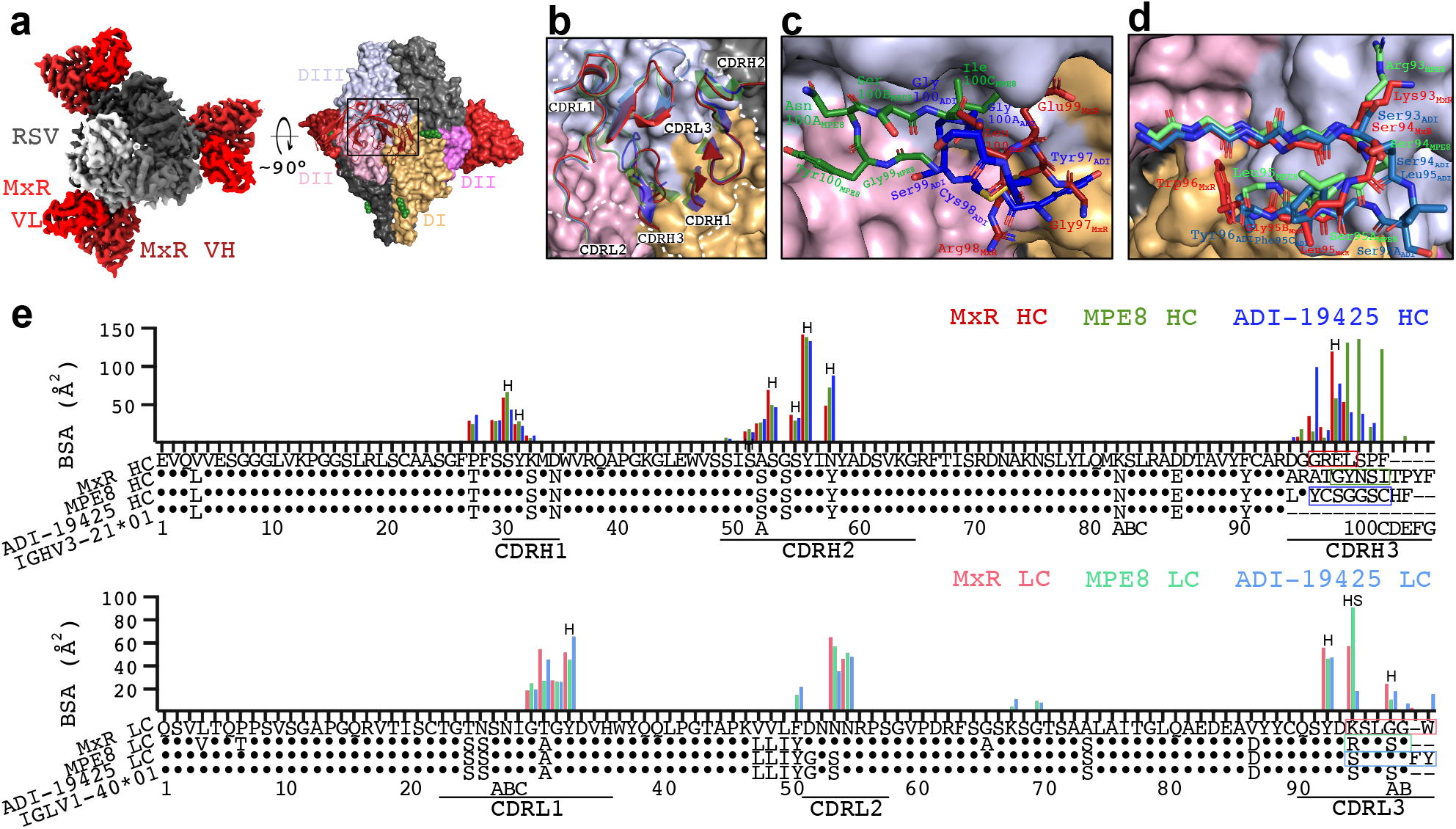
Cryo-EM structure of MxR Fab in complex with RSV preF and comparison with MPE8 and ADI-14925. **(a)** Left, a top-down view of the DeepEMhanced MxR:RSV cryo-EM map is shown, with MxR V_H_ shown in dark red, MxR V_L_ in light red, and RSV preF in shades of grey. Only the Fv domain of the F_ab_ is shown. Right, a surface representation of the structure is shown rotated 90 degrees. One preF protomer is colored per its structural domains, and the MxR Fv is shown in cartoon outline. The DII domain of the clockwise-adjacent protomer is in light pink, and designated DII’ to differentiate it from DII of the other protomer, colored in violet. Glycans are shown as green spheres. (**b**) Detailed view of the binding site on RSV preF with the CDRs of MxR, MPE8, and ADI-19425 superimposed and shown in cartoon. MPE8, in greens, and ADI-19425, in blues, are shown in transparency. Antigenic site III is designated by the white outline. (**c**) Zoomed in view of the CDRH3 binding site, rotated 45 degrees from panel (**b**). (**d**) Zoomed in view of the CDRL3 binding site, rotated 70 degrees clockwise from panel (**b**). The first four residues are shown as mainchain only to increase clarity and illustrate the similarity of CDRL3 between antibodies. (**e**) BSA plots for MxR, MPE8 and ADI-19425 residues which interact with RSV preF, atop a sequence alignment of the germline V-genes. The letter H indicates hydrogen bonds. The letter S indicates salt bridges. Residues in colored boxes correspond to the residues shown in panels (**b-d**).

The CDRH3 regions of these monoclonal antibodies have a greater degree of sequence and structural variation, with almost no conserved residues, and all bind near the DI/DII’/DIII interface within antigenic site III. There is a small cleft here, into which the CDRH3 loops can extend to varying degrees (**Fig. 4b**). MPE8 CDRH3 is the longest, while MxR is the shortest and ADI-19425 is of intermediate length. Notably, ADI-19425 uses a disulphide bond to stabilize this loop, whereas MPE8 and MxR lack this bond (**Fig. 4c**)^29, 31^. The rigid geometry imposed by the disulphide bond may impair ADI-19425’s ability to bind HMPV. The necessity of the MPE8 CDRH3 in forming the correct loop geometry to bind within the cleft of antigenic site III may thus be a structural basis behind MPE8’s reduced neutralization of HMPV compared to MxR (**Fig. 4b, c and Fig. 3d**). Due to a relatively shorter CDRH3 than MPE8, MxR binds at the DI/DIII interface, making no contact with DII’, and therefore does not require an extended loop to facilitate binding (**Fig. 4c and Fig. S5b**). Rather, only the CDRL2 of MxR contacts DII’ (**Fig. 4b**). The CDRL3 also shows some variation in interacting residues (**Fig. 4d**). For example, both MPE8 and MxR share a basic residue at position 93_LC_ which is not shared with ADI-19425, and all three antibodies exhibit slightly different loop arrangements in residues 94-96_LC_ (**Fig. 4d**). However, the binding mode of the CDRL3s appear broadly similar, without the unique structural features seen in the CDRH3.

### *In vivo* protection against viral infection

We next investigated whether the *in vitro* binding and neutralization data would translate into *in vivo* protection in an animal challenge model. Although the human parainfluenza viruses do not replicate in mice, upper and lower respiratory tract replication can be demonstrated in both hamsters and cotton rats^26, 32^. The hamster model has been used extensively to evaluate vaccine candidates to parainfluenza viruses, RSV, and HMPV^33, 34, 35, 36, 37^. Since all the viruses in this study could replicate in hamsters^33, 34, 38, 39, 40, 41, 42, 43^, we performed preclinical testing of MxR and 3×1 in the Golden Syrian hamster model. We performed intramuscular injections in hamsters at day -2, infected hamsters intranasally at day 0, and harvested lungs and nasal turbinates at day +5 post-infection to assess the efficacy of monoclonal antibodies as prophylaxis against infection (**Fig. 5a**). This dosing, route, and timing of administration are similar to those used in cotton rat models of RSV infection^44, 45, 46^. Prophylactic administration of 3×1 substantially reduced HPIV3 replication in the lungs by over 820-fold but had little impact on replication in the nasal turbinates (**Fig. 5b**). HPIV1 replication was completely blocked in the lungs of all but one animal and was significantly reduced in the nasal turbinates of hamsters that received 3×1 prophylaxis (**Fig. 5c**). Prophylactic administration of MxR completely suppressed RSV replication in the lungs and significantly reduced RSV replication in the nasal turbinates by over 34-fold (**Fig. 5d**). HMPV replication was significantly reduced by over 206-fold in the lungs of hamsters that received MxR prophylaxis (**Fig. 5e**).

**Figure 5.**
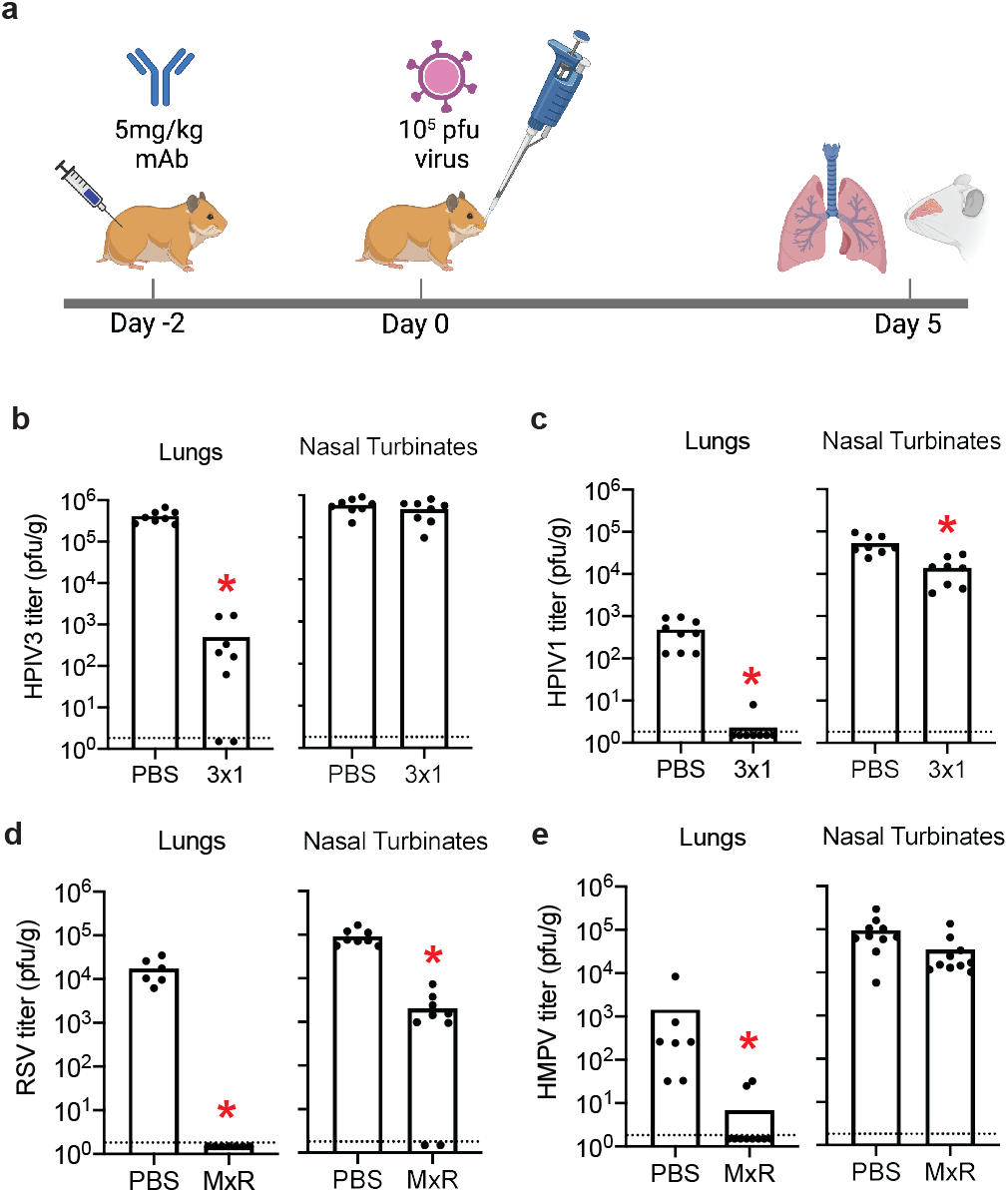
Efficacy of prophylactic administration of cross-neutralizing monoclonal antibodies 3×1 and MxR *in vivo*. **(a)** Schematic of experiments in which hamsters were injected intramuscularly with 5 mg/kg of 3×1 or MxR two days prior to intranasal challenge with 10^5^ pfu of HPIV3 (**b**), HPIV1 (**c**), RSV (**d**), or HMPV (**e**). Viral titers were measured by plaque assay in lung and nasal homogenates from individual mice (n=8-10) pooled from two independent experiments five days post-infection. Dashed lines indicate the limit of detection. Bars represent the mean, and asterisks indicate *p* < 0.01 by Mann-Whitney test compared to control hamsters injected with 1× DPBS.

If administered together as a cocktail, MxR and 3×1 could provide broad protection against HMPV, RSV, HPIV3, and HPIV1. Administration of a cocktail could also be useful in the setting of co-infections with multiple respiratory viruses, since co-infections can be associated with poorer outcomes in immunocompromised patients^16^. We therefore developed a co-infection model in hamsters to assess efficacy when MxR and 3×1 are co-administered together. We injected hamsters intramuscularly with a cocktail of MxR and 3×1 on day -2, co-infected hamsters with HPIV3 and RSV on day 0, and harvested lungs and nasal turbinates on day +5 post-infection (**Fig. 6a**). The cocktail of antibodies significantly reduced combined viral replication of HPIV3 and RSV in the lungs to undetectable levels in 6 out of 7 animals and by over 6-fold in nasal turbinates (**Fig. 6b, c**). Since the plaque assay to determine viral titers could not distinguish between HPIV3 vs. RSV, we developed custom TaqMan probes to quantify the individual viral loads by real-time PCR. The cocktail of antibodies did not have an impact on HPIV3 replication in the nasal turbinates but substantially reduced the viral load in the lungs by over 88-fold (**Fig. 6d, e**). The cocktail of antibodies specifically reduced RSV viral load in the lungs and nasal turbinates by over 17-fold and 2.9-fold, respectively, and RSV was below the limit of detection in the lungs of four out of seven hamsters (**Fig. 6f, g**).

**Figure 6.**
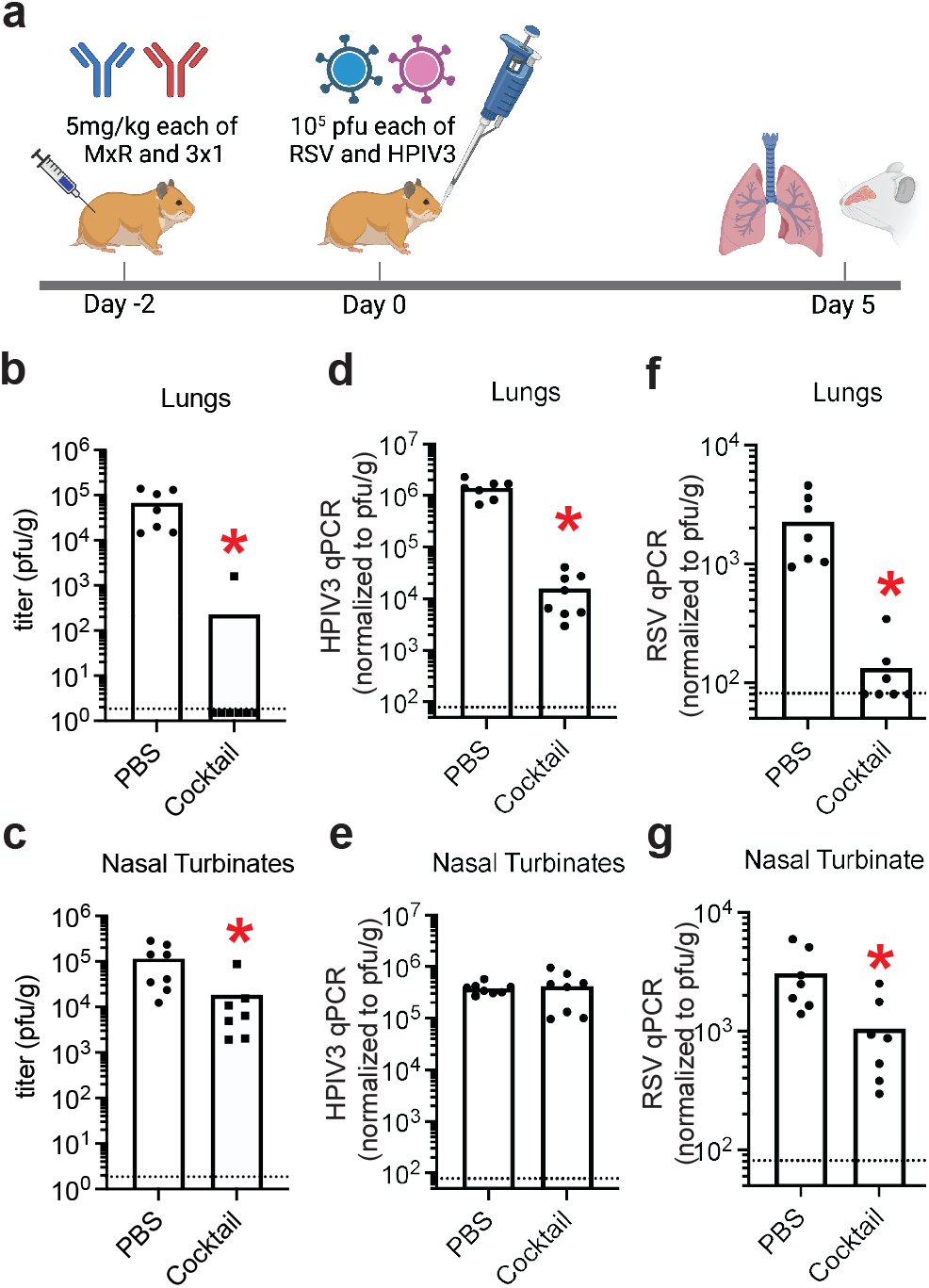
Efficacy of prophylactic administration of a cocktail 3×1 and MxR to protect against HPIV3 and RSV co-infection *in vivo*. (**a**) Schematic of two independent experiments in which hamsters (n=7-8) were injected intramuscularly with 5 mg/kg of 3×1 and MxR cocktail two days prior to intranasal challenge with 1×10^5^ pfu each of HPIV3 and RSV. Overall viral titer by plaque assay in lung (**b**) and nasal (**c**) homogenates at day five post-infection. The specific viral load of HPIV3 in lungs (**d**) and nasal (**e**) tissue homogenates was determined by real-time PCR using HPIV3-specific primers. The specific viral load of RSV in lungs (**f**) and nasal (**g**) tissue homogenates was determined by real-time PCR using RSV-specific primers. Dashed lines indicate the limit of detection, bars represent the mean, and asterisks denote *p* < 0.01 by Mann-Whitney test compared to control hamsters injected with 1× DPBS.

## DISCUSSION

We have isolated two anti-viral cross-neutralizing monoclonal antibodies: 3×1 which targets HPIV3 and HPIV1; and MxR which targets RSV and HMPV. Combined, these two antibodies could provide simultaneous and broad protection against four of the major respiratory viruses that afflict hematopoietic stem cell transplant patients and other vulnerable populations. We developed a “bait and switch” strategy based on the rationale that B cells capable of binding to one virus, while neutralizing another virus, are more likely to cross-neutralize both viruses. This strategy led to the discovery of 3×1, the first monoclonal antibody described that neutralizes multiple parainfluenza viruses. The “bait and switch” strategy could also be a generally useful approach for identifying cross-neutralizing antibodies against other pathogens in a situation where not all antigens are known or available. We also used magnetic enrichment and cell sorting to isolate rare B cells that could bind specifically to recombinant fusion proteins in the preF but not the postF conformation. Further, we leveraged feeder cells expressing CD40L, IL2, and IL21^27, 47^ to stimulate antibody production from individual B cells in culture. Together, these techniques permitted the high-throughput isolation and screening of B cells for cross-neutralization.

Targeting functionally conserved epitopes between homologous viruses is an attractive strategy to reduce the risk of developing drug resistance due to the emergence of escape mutations; and, at the same time, can increase the benefit of pre-exposure prophylaxis by protecting against a broader array of pathogens. The cross-neutralizing 3×1 mAb binds a novel site on HPIV3 preF, which we called site X, located between the equator and apex, on the vertices of the prefusion F trimer. It is possible that 3×1 binds the furin cleavage site of HPIV3, specifically the C-terminus of the F2 protein, although this region was too disordered to model effectively. Antibodies that cross-neutralize phylogenetically related viruses tend to target well-conserved epitopes^48^. The interaction between non-polar protrusions on the heavy chain of 3×1 with grooves on the surface could represent one mechanism that allows binding to both HPIV3 and HPIV1. Further, the light chain of 3×1 encompasses a large protrusion on HPIV3 preF and forms a hydrogen bond at Asp143, which is conserved in HPIV1. Further structural analysis will be needed to develop a better understanding of the molecular interactions that mediate 3×1 neutralization of HPIV1.

The MxR mAb neutralizes both HMPV and RSV and shares notable similarities with another antibody, MPE8^29, 49^, including significant sequence similarity that leads to comparable modes of binding. However, MxR is a more somatically hypermutated antibody and facilitates site III recognition without the extended CDRH3 seen in MPE8, which could provide more energetically favorable site recognition. This structural difference may be the basis for the observed difference between MxR and MPE8 in their neutralization of HMPV, with MxR having approximately 5.7-fold greater potency.

Palivizumab is currently the only FDA-approved antibody for prevention of RSV in high-risk infants. However, because the protection afforded by palivizumab is restricted to RSV, it has not gained widespread use for immunocompromised children or adults in whom other viruses like HMPV, HPIV3, and HPIV1 contribute significantly to disease^50^. A cocktail of MxR and 3×1 could therefore potentially fulfil this unmet need for a more broadly protective drug. The clinical indication for the RSV-specific D25 monoclonal antibody, which is being developed as nirsevimab to replace palivizumab, is similarly focused on infants^51^. We and others have observed reduced binding of D25 to the preF protein of RSV subtype B compared to RSV subtype A [**Fig. 3c**; also ref. ^52^]. This is notable because escape mutations to D25 have been identified in infants who suffered a breakthrough infection with RSV subtype B after receiving nirsevimab^53^. In contrast to D25, the cross-neutralizing MxR antibody we describe in the present study binds exceptionally well to the preF proteins of both RSV subtypes A and B.

To investigate the potential clinical utility of administering a cocktail of MxR and 3×1 to protect against RSV, HMPV, HPIV3, and HPIV1, we focused our *in vivo* efficacy studies on immunoprophylaxis. The importance of preventing respiratory viral infections for vulnerable populations has become increasingly apparent during the COVID-19 pandemic. A cocktail of two SARS-CoV-2-specific monoclonal antibodies marketed as Evusheld is now authorized by the FDA for prophylaxis in immunocompromised individuals who are expected to mount a poor response to vaccination. In a phase III trial, Evusheld, administered intramuscularly as 600 mg of total antibody, led to an 83% relative risk reduction in symptomatic COVID-19^54^. Due to escape mutations present in the omicron variant, the FDA revised the dose to 1200 mg of total antibody which, for a 60 kg individual, is a 20 mg/kg dose. In our *in vivo* efficacy studies, we similarly administered a cocktail of two antibodies MxR and 3×1, each at 5 mg/kg for a total 10 mg/kg dose. Therefore, the doses tested in the present study are within the range of other antibodies already in clinical use, leaving room for increased dosing in future human studies. Together, MxR and 3×1 represent promising mAb candidates for further development to protect against a broad array of respiratory viral infections in highly vulnerable patient populations.

## METHODS

### Study design

Peripheral blood was obtained by venipuncture from healthy, HIV-seronegative adult volunteers enrolled in the Seattle Area Control study, which was approved by the Fred Hutchinson Cancer Research Center institutional review board (Protocol #5567). PBMCs were isolated from whole blood using Accuspin System Histopaque-1077 (Sigma-Aldrich, cat#10771). Studies involving human spleens were deemed non-human subjects research since tissue was de-identified. Tissue fragments were passed through a basket screen, centrifuged at 300 × g for 7 minutes, incubated with ACK lysis buffer (Thermo Fisher, cat#A1049201) for 3.5 minutes, resuspended in RPMI (Gibco, cat#11875093), and passed through a stacked 500 µm and 70 µm cell strainer. Cells were resuspended in 10% dimethylsulfoxide in heat-inactivated fetal calf serum (Gibco, cat#16000044) and cryopreserved in liquid nitrogen before use.

### Cell lines

293F cells (Thermo Fisher, cat#R79007) were cultured in Freestyle 293 media (Thermo Fisher, cat#12338026). Vero cells (ATCC CCL-81), LLC-MK2 cells (ATCC CCL-7.1), and HEp-2 (ATCC CCL-23) were cultured in DMEM (Gibco, cat#12430054) supplemented with 10% fetal bovine serum and 100 U/ml penicillin plus 100 μg/mL streptomycin (Gibco, cat#15140122).

### Viruses

The recombinant viruses RSV-GFP, HMPV-GFP, HPIV1-GFP and HPIV3-GFP have been previously described^35, 55, 56, 57^ and were modified respectively from RSV strain A2 (GenBank accession number KT992094), HMPV CAN97-83 (GenBank accession number AY297749), HPIV1/Washington/20993/1964 (GenBank accession number AF457102), and HPIV3 JS (GenBank accession number Z11575) to express enhanced GFP. RSV subtype B strain 18537 (GenBank accession number MG813995) was obtained from ATCC (cat#VR-1580). Sendai virus strain Cantell (GenBank accession number AB855653) was obtained from ATCC (cat#VR-907). HMPV, HPIV1, and HPIV3 were cultured on LLC-MK2 cells, and RSV was cultured on HEp-2 cells. Virus was purified by centrifugation in a discontinuous 30%/60% sucrose gradient with 0.05 M HEPES and 0.1 M MgSO_4_ (Sigma-Aldrich, cat#H4034 and 230391, respectively) at 120,000 × g for 90 min at 4°C. Virus titers were determined by infecting Vero cell monolayers in 24-well plates with serial 10-fold dilutions of virus, overlaying with DMEM containing 4% methylcellulose (Sigma-Aldrich, cat#M0387). For assays involving HPIV1, HMPV, and Sendai virus, 1.2% of 0.05% of Trypsin (Gibco, cat#25300054) was included in the media. Fluorescent plaques were counted using a Typhoon scanner (GE Life Sciences) at five days post-infection.

### Expression and purification of antigens

Expression plasmids for His-tagged RSV, HMPV, and HPIV3 preF and postF antigens are previously described^28, 58, 59^. 293F cells were transfected at a density of 10^6^ cells/mL in Freestyle 293 media using 1 mg/mL PEI Max (Polysciences, cat#24765). Transfected cells were cultured for 7 days with gentle shaking at 37°C. Supernatant was collected by centrifuging cultures at 2,500 × g for 30 minutes followed by filtration through a 0.2 µM filter. The clarified supernatant was incubated with Ni Sepharose beads overnight at 4°C, followed by washing with wash buffer containing 50 mM Tris, 300 mM NaCl, and 8 mM imidazole. His-tagged protein was eluted with an elution buffer containing 25 mM Tris, 150 mM NaCl, and 500 mM imidazole. The purified protein was run over a 10/300 Superose 6 size exclusion column (GE Life Sciences, cat#17–5172–01). Fractions containing the trimeric F proteins were pooled and concentrated by centrifugation in a 50 kDa Amicon ultrafiltration unit (Millipore, cat#UFC805024). The concentrated sample was stored in 50% glycerol at −20°C.

### Tetramerization of antigens

Purified F antigens were biotinylated using an EZ-link Sulfo-NHS -LC-Biotinylation kit (Thermo Fisher, cat#A39257) with a 1:1.3 molar ratio of biotin to F. Unconjugated biotin was removed by centrifugation using a 50 kDa Amicon Ultra size exclusion column (Millipore). To determine the average number of biotin molecules bound to each molecule of F, streptavidin-PE (ProZyme, cat#PJRS25) was titrated into a fixed amount of biotinylated F at increasing concentrations and incubated at room temperature for 30 minutes. Samples were run on an SDS-PAGE gel (Invitrogen, cat#NW04127BOX), transferred to nitrocellulose, and incubated with streptavidin–Alexa Fluor 680 (Thermo Fisher, cat#S32358) at a dilution of 1:10,000 to determine the point at which there was excess biotin available for the streptavidin–Alexa Fluor 680 reagent to bind. Biotinylated F was mixed with streptavidin-allophycocyanin (APC) or streptavidin-PE at the ratio determined above to fully saturate streptavidin and incubated for 30 minutes at room temperature. Unconjugated F was removed by centrifugation using a 300 K Nanosep centrifugal device (Pall Corporation, cat#OD300C33). APC/DyLight755 and PE/DyLight650 tetramers were created by mixing F with streptavidin-APC pre–conjugated with DyLight755 (Thermo Fisher, cat#62279) or streptavidin-PE pre-conjugated with DyLight650 (Thermo Fisher, cat# 62266), respectively, following the manufacturer’s instructions. On average, APC/DyLight755 and PE/DyLight650 contained 4–8 DyLight molecules per APC and PE. The concentration of each tetramer was calculated by measuring the absorbance of APC (650 nm, extinction coefficient = 0.6 µM^−1^ cm^−1^) or PE (566 nm, extinction coefficient = 2.0 µM^−1^ cm^−1^).

### Tetramer enrichment

1–2 × 10^8^ frozen PBMCs or 4–8 × 10^7^ frozen spleen cells were thawed into DMEM with 10% fetal calf serum and 100 U/ml penicillin plus 100 µg/ml streptomycin. Cells were centrifuged and resuspended 50 µL of ice-cold fluorescence-activated cell sorting (FACS) buffer composed of phosphate-buffered saline (PBS) and 1% newborn calf serum (Thermo Fisher, cat#26010074). PostF APC/DyLight755 and PE/Dylight650 conjugated tetramers were added at a final concentration of 25 nM in the presence of 2% rat and mouse serum (Thermo Fisher) and incubated at room temperature for 10 min. PreF APC and PE tetramers were then added at a final concentration of 5 nM and incubated on ice for 25 min, followed by a 10 mL wash with ice-cold FACS buffer. Next, 50 μL each of anti-APC-conjugated (Miltenyi Biotec, cat#130–090– 855) and anti-PE-conjugated (Miltenyi Biotec, cat#130-048-801) microbeads were added and incubated on ice for 30 min, after which 3 mL of FACS buffer was added and the mixture was passed over a magnetized LS column (Miltenyi Biotec, cat#130–042– 401). The column was washed once with 5 mL ice-cold FACS buffer and then removed from the magnetic field and 5 mL ice cold FACS buffer was pushed through the unmagnetized column twice using a plunger to elute the bound cell fraction.

### Flow cytometry

Cells were incubated in 50 μL of FACS buffer containing a cocktail of antibodies for 30 minutes on ice prior to washing and analysis on a FACS Aria (BD). Antibodies included anti-IgM FITC (G20–127, BD), anti-CD19 BUV395 (SJ25C1, BD), anti-CD3 BV711 (UCHT1, BD), anti-CD14 BV711 (M0P-9, BD), anti-CD16 BV711 (3G8, BD), anti-CD20 BUV737 (2H7, BD), anti-IgD BV605 (IA6–2, BD), anti-CD27 PE/Cy7 (LG.7F9, eBioscience), and a fixable viability dye (Tonbo Biosciences, cat#13–0870– T500). B cells were individually sorted into either 1) empty 96-well PCR plates and immediately frozen, or 2) flat bottom 96-well plates containing feeder cells that had been seeded at a density of 28,600 cells/well one day prior in 100 µL of IMDM media (Gibco, cat#31980030) containing 10% fetal calf serum, 100 U/ml penicillin plus 100 µg/ml streptomycin, and 2.5 µg/mL amphotericin. B cells sorted onto feeder cells were cultured at 37°C for 13 days.

### B cell receptor sequencing

For individual B cells sorted and frozen into empty 96-well PCR plates, reverse transcription (RT) was directly performed after thawing plates using SuperScript IV (Thermo Fisher, cat#18090200) as previously described^60, 61^. Briefly, 3 µL RT reaction mix consisting of 3 µL of 50 µM random hexamers (Thermo Fisher, cat#48190011), 0.8 µL of 25 mM deoxyribonucleotide triphosphates (dNTPs; Thermo Fisher, cat#N8080261), 1 µL (20 U) SuperScript IV RT, 0.5 µL (20 U) RNaseOUT (Thermo Fisher, cat#10777019), 0.6 µL of 10% Igepal (Sigma-Aldrich, cat#I8896), and 15 µL RNase-free water was added to each well containing a single sorted B cell and incubated at 50°C for 1 hour. For individual B cells sorted onto feeder cells, supernatant was removed after 13 days of culture, plates were immediately frozen on dry ice, stored at −80°C, thawed, and RNA was extracted using the RNeasy Micro Kit (Qiagen, cat#74034). The entire eluate from the RNA extraction was used instead of water in the RT reaction. Following RT, 2 µL of cDNA was added to 19 µl PCR reaction mix so that the final reaction contained 0.2 µL (0.5 U) HotStarTaq Polymerase (Qiagen, cat#203607), 0.075 µL of 50 µM 3′ reverse primers, 0.115 µL of 50 µM 5′ forward primers, 0.24 µL of 25 mM dNTPs, 1.9 µL of 10× buffer (Qiagen), and 16.5 µL of water. The PCR program was 50 cycles of 94°C for 30 s, 57°C for 30 s, and 72°C for 55 s, followed by 72°C for 10 min for heavy and kappa light chains. The PCR program was 50 cycles of 94°C for 30 s, 60°C for 30 s, and 72°C for 55 s, followed by 72°C for 10 min for lambda light chains. After the first round of PCR, 2 µL of the PCR product was added to 19 µL of the second-round PCR reaction so that the final reaction contained 0.2 µL (0.5 U) HotStarTaq Polymerase, 0.075 µL of 50 µM 3′ reverse primers, 0.075 µL of 50 µM 5′ forward primers, 0.24 µL of 25 mM dNTPs, 1.9 µL 10× buffer, and 16.5 µL of water. PCR programs were the same as the first round of PCR. 4 μL of the PCR product was run on an agarose gel to confirm the presence of a ∼500-bp heavy chain band or 450-bp light chain band. 5 μL from the PCR reactions showing the presence of heavy or light chain amplicons was mixed with 2 µL of ExoSAP-IT (Thermo Fisher, cat#78201) and incubated at 37°C for 15 min followed by 80° C for 15 min to hydrolyze excess primers and nucleotides. Hydrolyzed second-round PCR products were sequenced by Genewiz with the respective reverse primer used in the second round PCR, and sequences were analyzed using IMGT/V-Quest to identify V, D, and J gene segments. Paired heavy chain VDJ and light chain VJ sequences were cloned into pTT3-derived expression vectors containing the human IgG1, IgK, or IgL constant regions using In-Fusion cloning (Clontech, cat#638911) as previously described^62^.

### Monoclonal antibody production

Secretory IgG was produced by co-transfecting 293 F cells at a density of 10^6^ cells/mL with the paired heavy and light chain expression plasmids at a ratio of 1:1 in Freestyle 293 media using 1 mg/mL PEI Max. Transfected cells were cultured for 7 days with gentle shaking at 37°C. Supernatant was collected by centrifuging cultures at 2,500 × g for 15 minutes followed by filtration through a 0.2 µM filter. Clarified supernatants were then incubated with Protein A agarose (Thermo Scientific, cat#22812) followed by washing with IgG binding buffer (Thermo Scientific, cat#21007). Antibodies were eluted with IgG Elution Buffer (Thermo Scientific, cat#21004) into a neutralization buffer containing 1 M Tris-base pH 9.0. Purified antibody was concentrated and buffer exchanged into PBS using an Amicon ultrafiltration unit with a 50 kDa molecular weight cutoff.

### Bio-layer interferometry

Bio-layer interferometry (BLI) assays were performed on the Octet.Red instrument (ForteBio) at room temperature with shaking at 500 rpm. For apparent affinity (*K*_D_) analyses, anti-penta His capture sensors (ForteBio, cat#18–5120) were loaded in kinetics buffer (PBS with 0.01% bovine serum albumin, 0.02% Tween 20, and 0.005% NaN_3_, pH 7.4) containing 0.5 µM purified RSV-A, RSV-B, HMPV, or HPIV3 preF for 150 s. After loading, the baseline signal was recorded for 60 s in kinetics buffer. The sensors were then immersed in kinetics buffer containing 266.7, 133.3, 66.7, 33.3, or 16.7 nM of purified monoclonal antibody for a 300 s association step followed by immersion in kinetics buffer for a dissociation phase of at least 600 s. The maximum response was determined by averaging the nanometer shift over the last 5 s of the association step after subtracting the background signal from each analyte-containing well using a negative control mAb at each time point. Curve fitting was performed using a 1:1 binding model and ForteBio data analysis software. For competitive binding assays, anti-penta His capture sensors were loaded in kinetics buffer containing 1 µM His-tagged HPIV3 preF for 300 s. After loading, the baseline signal was recorded for 30 s in kinetics buffer. The sensors were then immersed for 300 s in kinetics buffer containing 40 µg/mL of the first antibody followed by immersion for another 300 s in kinetics buffer containing 40 µg/mL of the second antibody. Percent competition was determined by dividing the maximum increase in signal of the second antibody in the presence of the first antibody by the maximum signal of the second antibody alone.

### Neutralization assays

For neutralization screening of culture supernatants, Vero cells were seeded in 96-well flat bottom plates and cultured for 48 hours. After 13 days of culture, 40 µL of B cell culture supernatant was mixed with 25 µL of sucrose-purified GFP-HPIV1 diluted to 2,000 plaque forming units (pfu)/mL for 1 hour at 37°C. Vero cells were then incubated with 50 µL of the supernatant/virus mixture for 1 hour at 37°C to allow viral adsorption. Next, each well was overlaid with 100 µL DMEM containing 4% methylcellulose and 1.2% of 0.05% trypsin. Fluorescent plaques were counted at 5 days post-infection using a Typhoon imager.

Neutralizing titers of monoclonal antibodies were determined by a 60% plaque reduction neutralization test (PRNT_60_). Vero cells were seeded in 24-well plates and cultured for 48 hours. Monoclonal antibodies were serially diluted 1:4 in 120 µL DMEM and mixed with 120 µL of sucrose-purified RSV, HMPV, HPIV3, or HPIV1 diluted to 2,000 pfu/mL for one hour at 37°C. Vero cells were incubated with 100 µL of the antibody/virus mixture for 1 hour at 37°C to allow viral adsorption. Each well was then overlaid with 500 µL DMEM containing 4% methylcellulose. Fluorescent plaques were counted at 5 days post-infection using a Typhoon imager. PRNT_60_ titers were calculated by linear regression analysis (http://exon.niaid.nih.gov/plaquereduction/).

### Cryo-EM complex and grid preparation

For MxR, 1.47 mg of RSV preF with 1.45 mg of MxR Fab overnight at 4 ºC with gentle rocking. Complexes were purified over a 10/300 Superose 6 column. A very broad peak eluted, and the nine earliest fractions containing appreciable amounts of protein were collected, pooled, and concentrated to 0.22 mg/mL using a 10 kDa cutoff Amicon ultrafiltration unit (Sigma-Aldrich, cat#UFC8030). For 3×1, we prepared complexes by incubating 50 µg of HPIV3 preF with 150 µg of 3×1 Fab overnight at 4 ºC with gentle rocking. The complexes were purified over a Superdex 200 16/600 SEC column (Cytiva, cat#28989335). A single narrow high molecular weight peak eluted and was collected, pooled, and concentrated to 0.4 mg/mL using a 10 kDa cutoff Amicon ultrafiltration unit.

Complexes were frozen on Quantifoil 1.2/1.3 UltrAu 300 mesh grids (Electron Microscopy Sciences, cat#Q350AR13A) using a Vitrobot Mk. IV (ThermoFisher). Dodecyl-β-d-maltoside (DDM) to a concentration of 0.05% was added 30 minutes prior to the start of freezing. Grids were also glow discharged prior to the start of freezing. Vitrobot conditions were kept at 22ºC and 100% humidity throughout grid preparation. Numerous variations to concentration, blot conditions, application, and time were used to prepare grids for screening. The final MxR:RSV grids used for collection were frozen with 4 µL of sample at 0.21 mg/mL, a 14 second blot time, 0 blot force, and a 5 second wait between application and blotting. The final 3×1:HPIV3 grids used for collection were frozen with 4 µL of sample at 0.19 mg/mL, a 12 second blot time, 0 blot force, and a 15 second wait between application and blotting.

### Data collection, processing, and model refinement

Grids were collected on a Krios 300kV Cryo-EM scope with a K3 camera at the Pacific Northwest Center for Cryo-EM (PNCC). Data was collected in 50 frame movies using Serial EM with a super resolution of 0.514425 Å/px and an exposure of 50 e-/Å^2^. Both collections ran for 24 hours, producing 6282 movies for 3×1:HPIV3 and 5796 movies for MxR:RSV. Notably, 3×1:HPIV3 was collected at a 30º tilt due to observed preferred orientation during screening.

Data sets were processed using cryoSPARC v3.3.1. Movies were imported, patch motion corrected, and underwent patch contrast transfer function (CTF) estimation. Micrographs were curated with the major criteria being CTF fitting < 4 Å. A small subset of particles (∼50,000) was picked, underwent 2D classification, and classes selected for the template-based picking job. Picks were inspected, curated, and extracted with 1.49 million for MxR:RSV and 2.02 million for 3×1:HPIV3, both at a box size of 192 px at ∼2.2 Å/px.

We performed two rounds of 2D classification with 100 classes each on the MxR:RSV particles, selecting for only particles displaying three Fabs and a well-ordered complex without artifacts. The output was 377,982 particles. A single ab-initio model was generated and refined, with a GSFSC resolution significantly greater than Nyquist at 2.0577 Å/px. Particles were re-extracted at 1.02885 Å/px and subject to local CTF refinement and non-uniform refinement with C3 symmetry. Following this, exposures were further curated and particles reextracted using the Local Motion Correction job and binned to 1.02885 Å/px, producing a stack of 354,958 particles. Non-uniform 3D refinement produced a sharpened map with a GSFSC resolution of 2.41 Å at the 0.143 cutoff using C1 symmetry and a resolution of 2.24 Å at the 0.143 cutoff using C3 symmetry and with a custom mask which cropped out the CH1 of each Fab and the trimerization domain on RSV preF. The unmasked C3 map has a resolution of 2.8 Å.

The template generation for 3×1:HPIV3 produced a mix of 1 Fab, 2 Fabs, and 3 Fabs, all of which were used for original template picking. We performed one round of 2D classification with 100 classes, selecting for solitary particles which showed any number of Fabs bound, producing a stack of 1.3 million particles. Ab-initio modelling produced a single map with one Fab bound to HPIV3 preF, and subsequent refinement using non-uniform refinement produced a map with a GSFSC resolution of 3.62 Å at the 0.143 cutoff using C1 symmetry. An additional three Fab map was produced with C3 symmetry in non-uniform refinement to 4.3 Å, though due to the low resolution this map was not used in model building. Minor improvements were made by performing a C3 symmetry homogenous refinement, followed by C3 symmetry expansion, 3D classification with four classes, and the removal of duplicate particles from the most homologous class, producing a stack of 1.04 million particles. Particles were re-extracted with the Local Motion Correction job, binned to 1.02885 Å/px, and the map refined using non-uniform refinement. This produced our final map with a resolution of 3.62 Å at the 0.143 cutoff by GSFSC.

Both maps were further processed using the COSMIC^2^ computer and the DeepEMhancer module. Both the DeepEM enhanced and cryoSPARC sharp maps were used in fitting the structures of MxR:RSV and 3×1:HPIV3 to the map, though only the original cryoSPARC sharp maps were used in the final real-space refinement in Phenix. Unmasked half maps, the sharp EM map, and the refine mask were deposited to the PDB and EMDB. Structure refinement was also carried out in ChimeraX using the ISOLDE plugin as necessary to fit low resolution regions. All graphics were produced in Pymol. Graphs were prepared using GraphPad Prism 9. Buried surface area analysis was carried out using the PDBePISA server.

### Animals and viral challenge

All procedures were reviewed and approved by the Fred Hutch Institutional Animal Care and Use Committee and conducted in accordance with institutional and National Institutes of Health guidelines. Golden Syrian hamsters were infected intranasally with 100 µL of 10^5^ pfu RSV, HMPV, HPIV3, or HPIV1 in two independent experiments. This sample size is consistent with previously published experiments testing the efficacy of RSV monoclonal antibodies in the cotton rat model^44, 45, 63, 64^. Monoclonal antibody was administered intramuscularly at 5 mg/kg in 50 µL PBS two days prior to infection. For the co-infection model, 10^5^ pfu of each virus was mixed in 100 µL 1× DPBS, and 5 mg/kg of each monoclonal antibody was mixed in 50 µL 1× DPBS. Nasal turbinates and lungs were removed for viral titration by plaque assay 5 days post-infection and clarified by centrifugation in DMEM. Confluent Vero cell monolayers were inoculated in duplicate with diluted homogenates in 24-well plates. After incubating for 1 hour at 37°C, wells were overlaid with 4% methylcellulose (made with 1.2% of 0.05% trypsin for specimens from animals challenged with HMPV and HPIV1). After five days, plaques were counted using the Typhoon imager to determine titers as pfu per gram of tissue. Aliquots of nasal turbinate and lung samples were also saved for quantification of viral load by real-time PCR.

### Real-time PCR

Viral RNA was extracted from 140 µL of sample homogenate using the QIAamp vRNA mini kit (Qiagen, cat#52904). RNA was eluted with 50 µL water and 11 µL of the eluate was used for reverse transcription. Custom reverse transcription primers for RSV (5’- TCCAGCAAATACACCATCCAAC-3’) and HPIV3 (5’- CTAGAAGGTCAAGAAAAGGGAACTC-3’) were designed to specifically bind to the genomes of RSV and HPIV3, respectively. One microliter of each primer at 2 µM was included in a RT reaction mix containing 1 µL of RNaseOut, 1 µL of 0.1 M DTT, 4 µL of SuperScript IV buffer, and 1 µL of SuperScript IV reverse transcriptase (Thermo Fisher, cat#18090200). Reverse transcription was performed with the following cycle: 42°C for 10 min, 20°C for 10 min, 50°C for 10 min, and 80°C for 10 min. Custom TaqMan Gene Expression Assays were developed for RSV (forward primer 5’-TGACTCTCCTGATTGTGGGATGATA-3’, reverse primer 5’- CGGCTGTAAGACCAGATCTGT-3’, and reporter 5’- CCCCTGCTGCTAATTT-3’) & HPIV3 (forward primer, 5’- CGGTGACACAGTGGATCAGATT-3’, reverse primer 5’- TGTTTCAACCATAAGAGTTACCAAGCT-3’, and reporter 5’- ACCGCATGATTGACCC-3’). The PCR reaction consisted of 2.5 µL of these primers, 10 µL of the reverse transcription reaction, 25 µL of TaqMan Universal Master Mix II with UNG (Thermo Fisher, cat#4440038), and 12.5 µL water. Real-time PCR was performed using the QuantStudio 7 Flex Real-Time PCR System with the following parameters: 50°C for 2 min and 95°C for 10 min followed by 40 cycles at 95 °C for 15 s and 60 °C for 1 min. To generate a standard curve, viral RNA was extracted from sucrose purified viral stocks of RSV with known titers in pfu/mL. Reverse transcription was performed as above. The reverse transcription reaction was serially diluted eight times at 1:4 in water. Real-time PCR was performed as above, and standard curves were generated to interpolate viral loads to pfu/g using QuantStudio Real-time PCR Software v1.0.

### Statistical analysis

Statistical analysis was performed using GraphPad Prism 7. Pairwise statistical comparisons were performed using Mann-Whitney two-tailed testing. *p* < 0.01 was considered statistically significant. Data points from individual samples are displayed.

## ACKNOWLEDGMENTS

We thank Julie McElrath for PBMCs from the Seattle Area Control cohort; Leo Stamatatos for the use of laboratory space and equipment; Ramasamy Bakthavatsalam and LifeCenter Northwest for providing de-identified spleen remnants; Ursula Buchholz, Shirin Munir, and Peter Collins for providing the GFP-expressing RSV, HMPV, HPIV3, and HPIV1; Peter Kwong, Guillaume Stewart-Jones, and Aliaksandr Druz for expression of HPIV3 preF; Barney Graham for expression of RSV preF; Theodore Jardetzky and Xiaolin Wen for expression of HMPV preF; Steve Voght for proof-reading the manuscript; Paula Culver, Francesca Urselli, and Laura Yates for administrative support; the Fred Hutchinson Flow Cytometry Shared Resources for assistance with instruments; the Fred Hutchinson Comparative Medicine Shared Resources for assistance with housing hamsters; and the Taylor lab for helpful discussions. Experimental schematics were created with BioRender.com.

## AUTHOR CONTRIBUTIONS

M.C. and M.B. designed and conducted the experiments, analyzed the data, and wrote the manuscript. M.D.G. conducted experiments, analyzed data, and edited the manuscript. J.V.R. and M.P coordinated and performed the structural analysis. J.T.T. conceived the study, designed experiments, analyzed the data, and edited the manuscript. J.B. conceived the study, designed and conducted the experiments, analyzed the data, and wrote the manuscript.

## DATA AVAILABILITY

Sequencing and structural data that support the findings of this study have been deposited in the Protein Data Bank (PDB) and Electron Microscopy Data Bank (EMDB) and are accessible through accession numbers PDB 8DG8 (EMDB 27418) for 3×1/HPIV3 and PDB 8DG9 (EMDB 27419) for MxR/RSV.

## SUPPLEMENTARY FIGURES AND TABLES

**Supplementary Figure 1.**
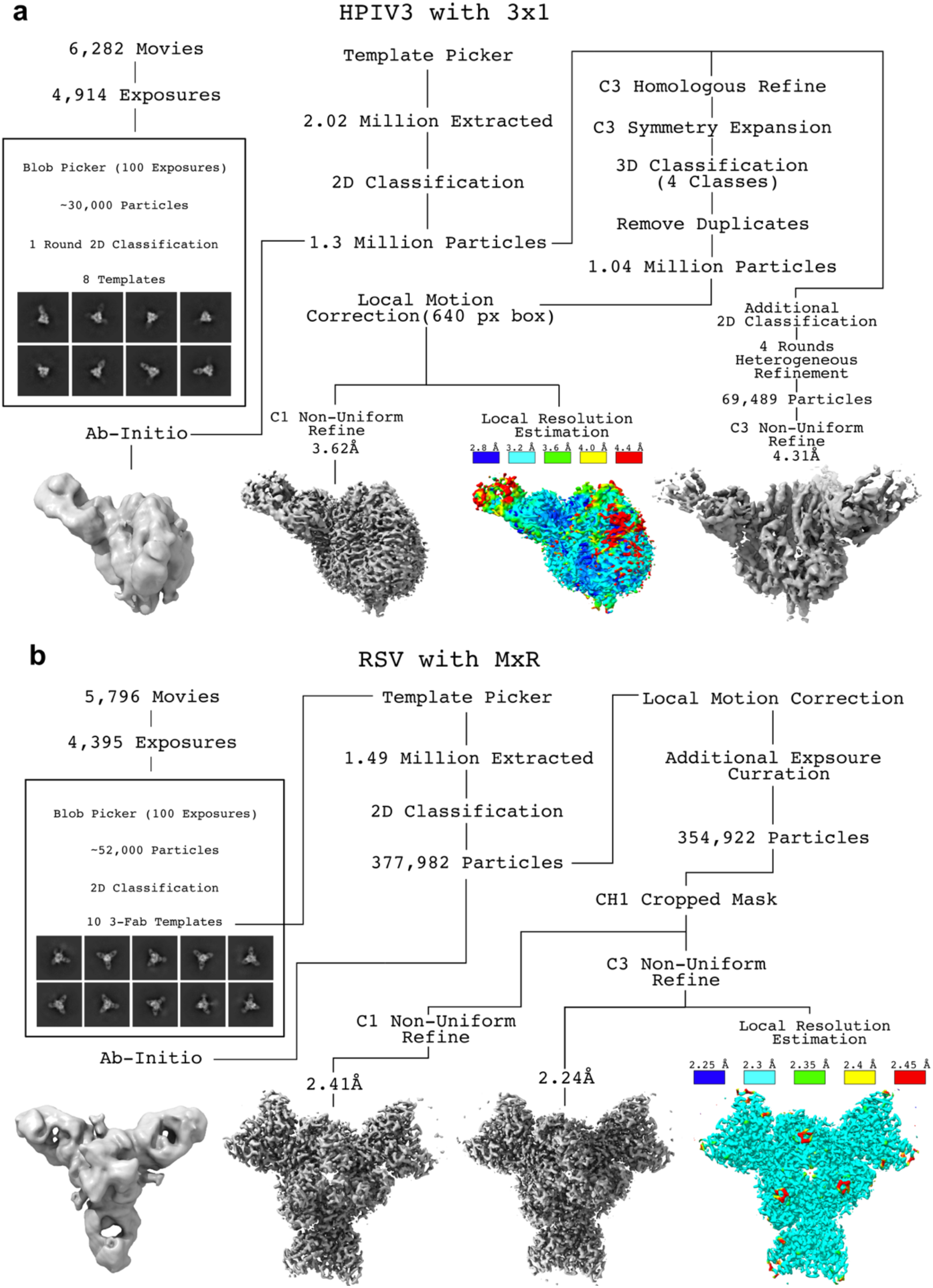
Cryo-EM map processing and resolution. (**a**) Overall processing pipeline for **Fig. 2** map refinement of the 3×1:HPIV3 complex. (**b**) Overall processing pipeline for **Fig. 4** map refinement of the MxR:RSV complex. Notable maps obtained during processing are shown in miniature. Template creation is shown and boxed, with templates used shown below, which double as representative 2D classes. Final maps are shown with local resolution estimation, with the color legend located above the map.

**Supplementary Figure 2.**
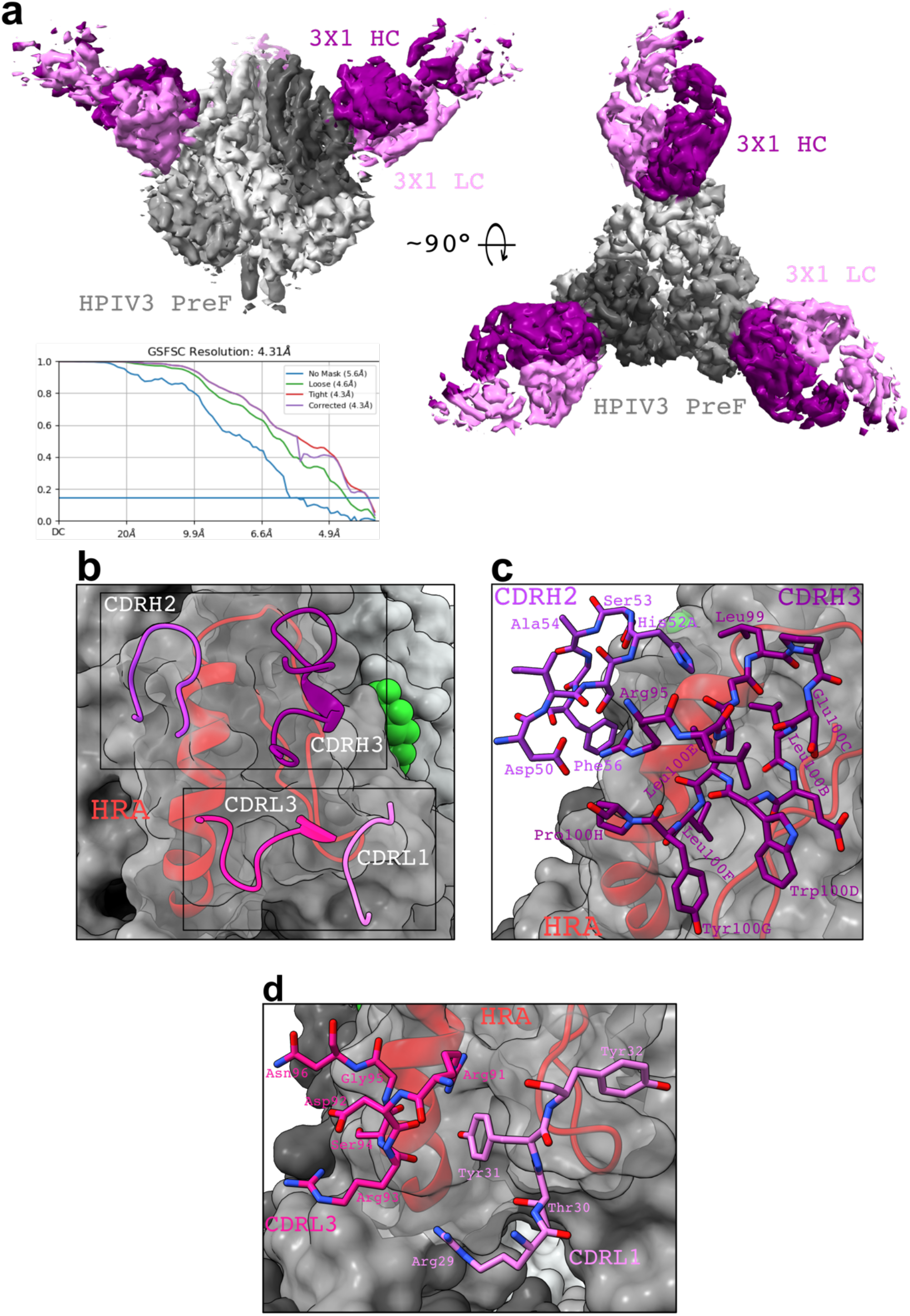
Three Fab structure of 3×1:HPIV3 and the HRA region that is bound by 3×1. (**a**) Top view and side view of a 4.31 Å structure of 3×1:HPIV3 obtained by C3 symmetry refinement. GS-FSC curve is shown below. (**b**) View of the 3×1 CDRs in contact with the HRA helix of HPIV3 preF. CDRs are labeled and colored distinctly, HRA is shown in red, and glycans are shown in green. (**c**) In-depth image of the CDRH2 and H3 bound to the HRA region. (**d**) In-depth image of the CDRL1 and L3 bound to the HRA region.

**Supplementary Figure 3.**
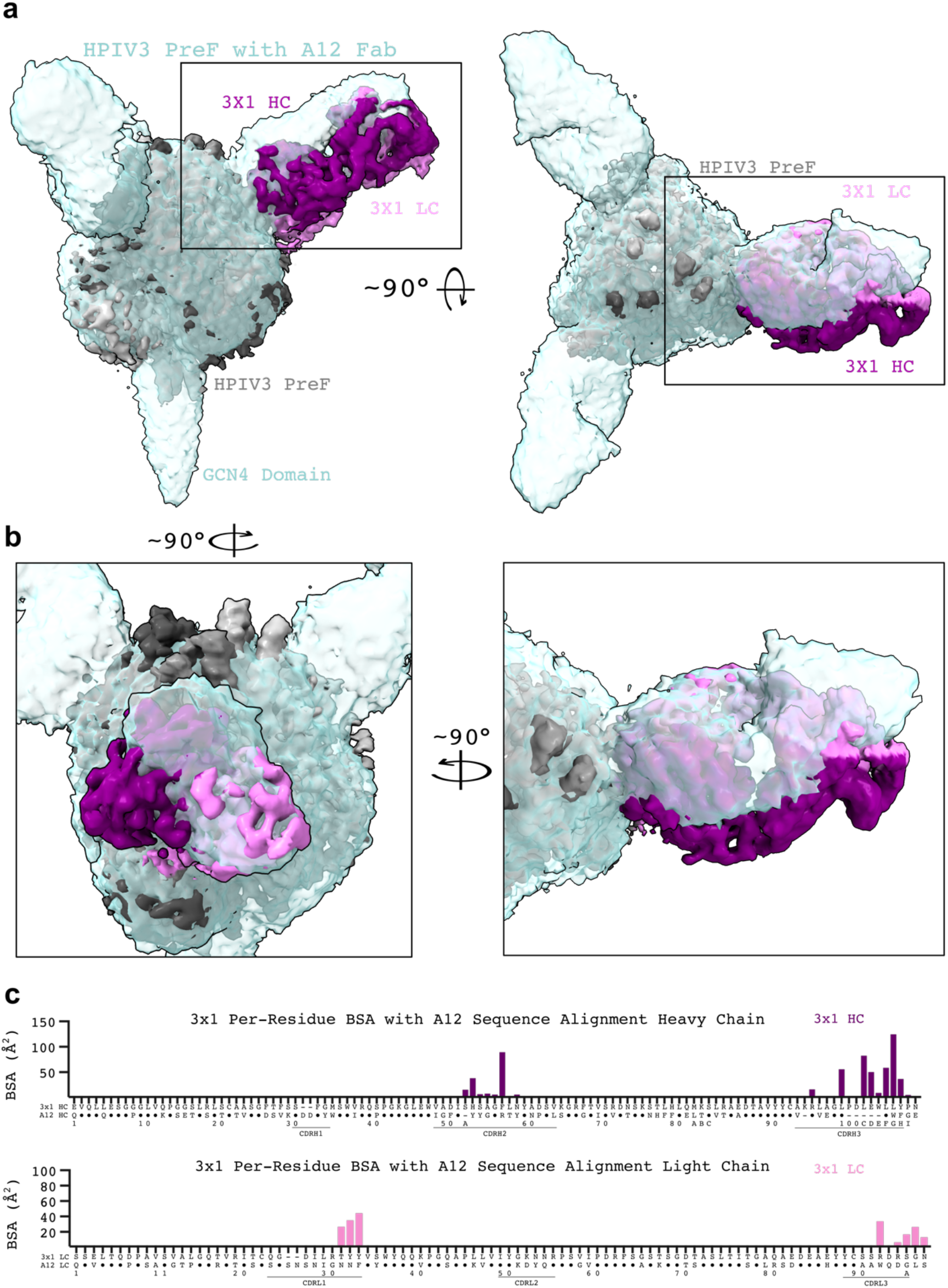
Comparison of 3×1:HPIV3 to negative stain electron microscopy (nsEM) data of PI3-A12:HPIV3. (**a**) Two views of the 3×1:HPIV3 cryo-EM map and PI3-A12:HPIV3 nsEM map are shown aligned. PI3-A12 is abbreviated A12. Labels show regions of interest in the corresponding color. Boxes indicate magnified area in panel (**b**). (**b**) Two zoomed in views show deviations in the binding orientation of 3×1 versus PI3-A12. (**c**) BSA plots show residues of 3×1 which interact with the HPIV3 preF protomer, atop a sequence alignment with PI3-A12.

**Supplementary Figure 4.**
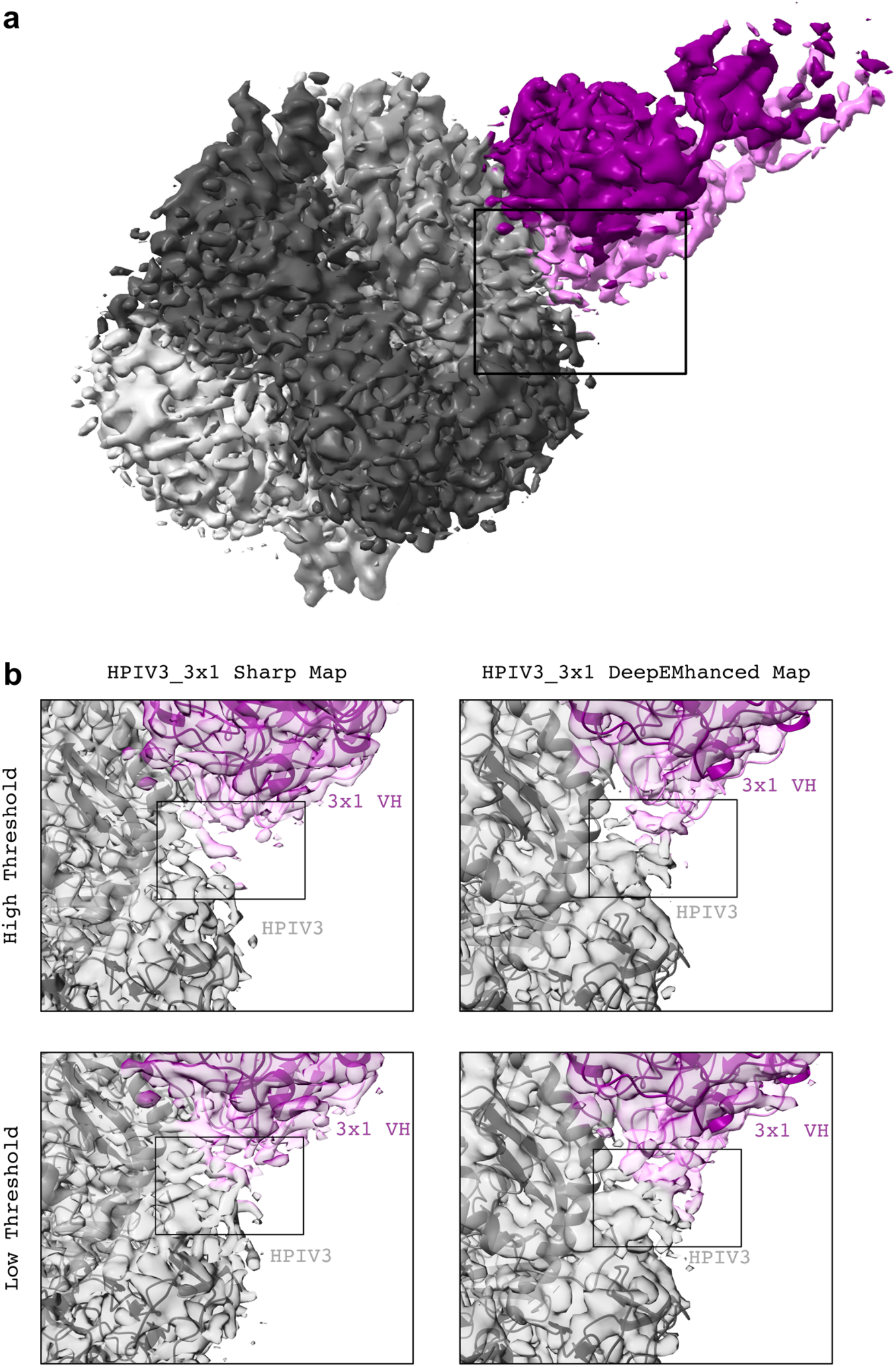
Putative HPIV3 furin cleavage site density maps. (**a**) Overall view of the 3×1:HPIV3 complex is shown with density corresponding to the cleavage site near unbound CDR residues on 3×1.**(b)** The region of interest is shown as both the sharp map and DeepEMhanced map at both high and low thresholds, with the high threshold approximately the normal structure building level, ∼5 sigma. The density has a resemblance to a sheet-turn-sheet motif which would be appropriate for the C-terminus of the F2 protein product following furin cleavage.

**Supplementary Figure 5.**
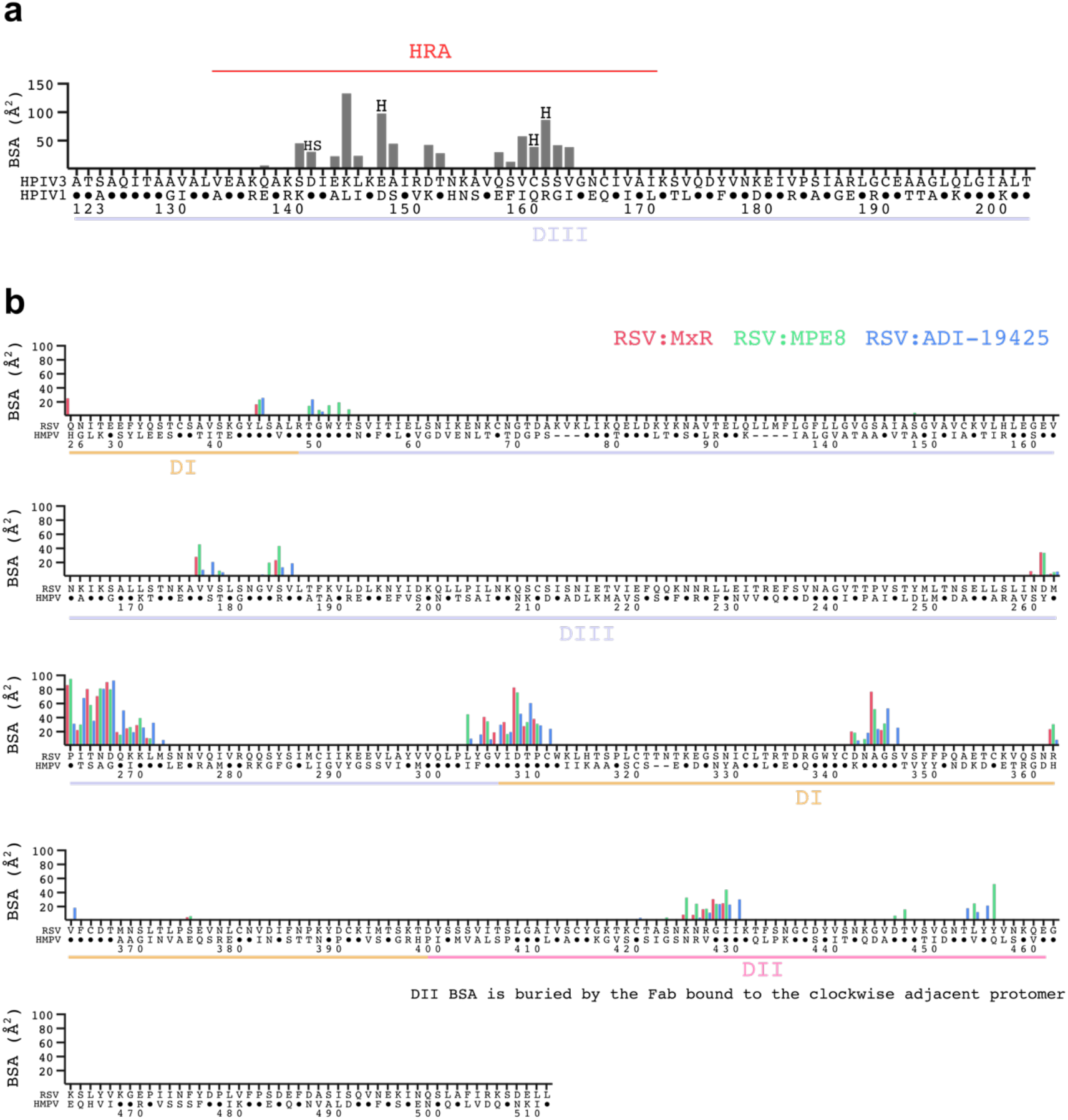
Per residue BSA of interacting RSV and HPIV3 preF residues. (**a**) BSA plot of HPIV3 residues which contact 3×1; this is a complementary plot to **Fig. 2f**. Only residues 123-204 – all within domain III – make contact with 3×1. Below the HPIV3 sequence, the HPIV1 sequence is aligned. (**b**) BSA plot showing RSV residue interactions with mAbs in MxR:RSV, MPE8:RSV, and ADI-19425:RSV; this data is complementary to **Fig. 4e**. The sequences of RSV and HMPV F are aligned below. Below that, colored bars indicate the structural domains of RSV using the same colors as **Fig. 4a**.

**Supplementary Figure 6.**
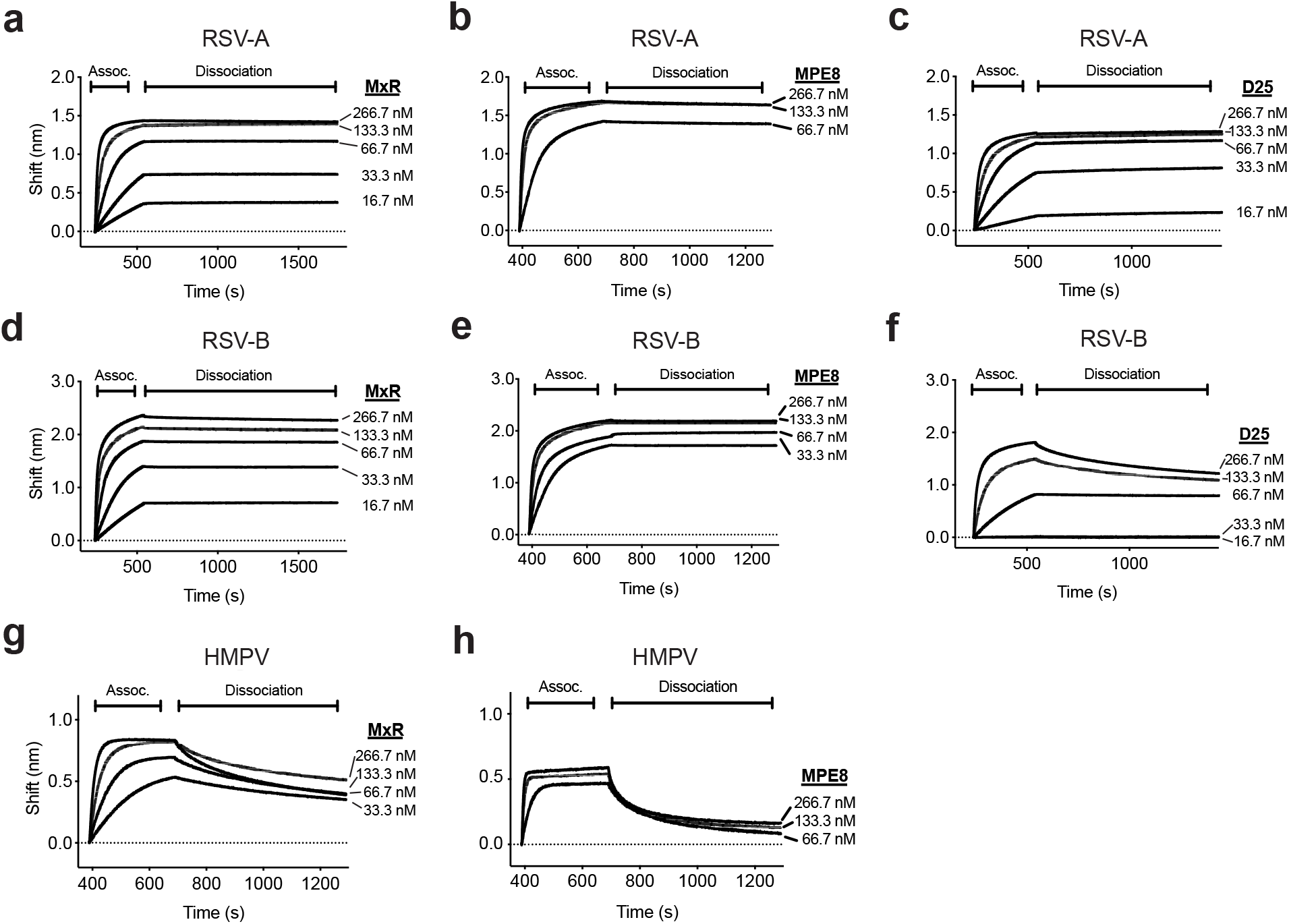
Binding kinetics of cross-neutralizing monoclonal antibodies. Apparent affinity (*K*_D_) of MxR (**a, d, g**), MPE8 (**b, e, h**), and D25 (**c, f**) at concentrations ranging from 16.7-266.7 nM to RSV-A (**a-c**), RSV-B (**d-f**), and HMPV (**g-h**) preF was determined by BLI. Penta-His capture sensors were loaded with His-tagged preF proteins. Association with MxR, MPE8, and D25 was then measured for 300 s followed by dissociation for at least 600 s. All measurements are normalized against an isotype control antibody.

**Supplementary Figure 7.**
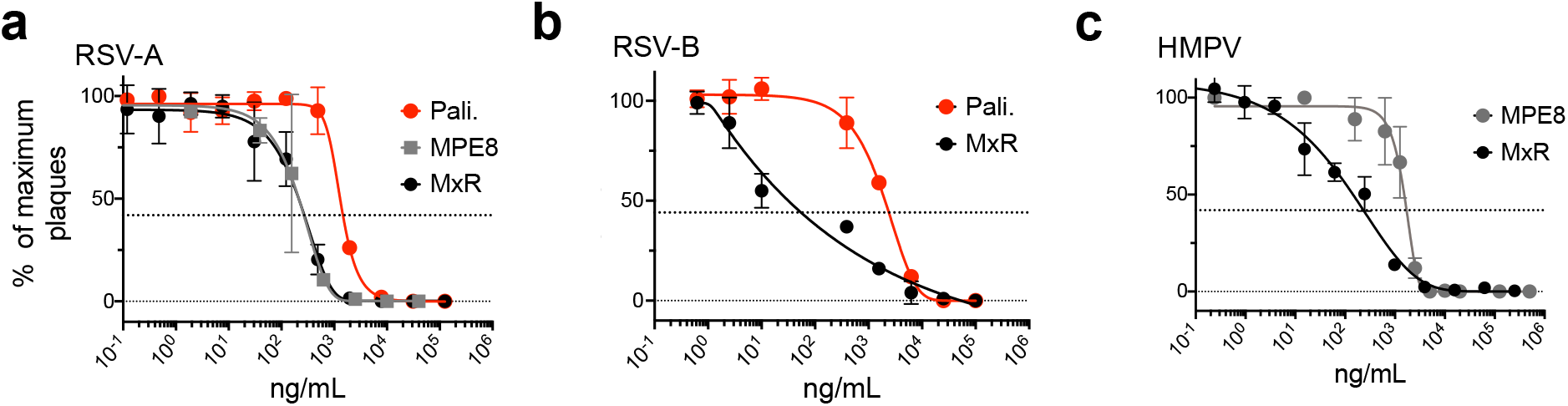
Neutralization potency *in vitro* of cross-neutralizing monoclonal antibodies. Vero cells were infected with RSV-A, RSV-B, or HMPV in the presence of serial dilutions of palivizumab (abbreviated Pali.), MPE8, or MxR. The dotted midline indicates the PRNT_60_. Data points are from three independent experiments with each experiment consisting of two technical replicates.

**Supplemental Table 1.**
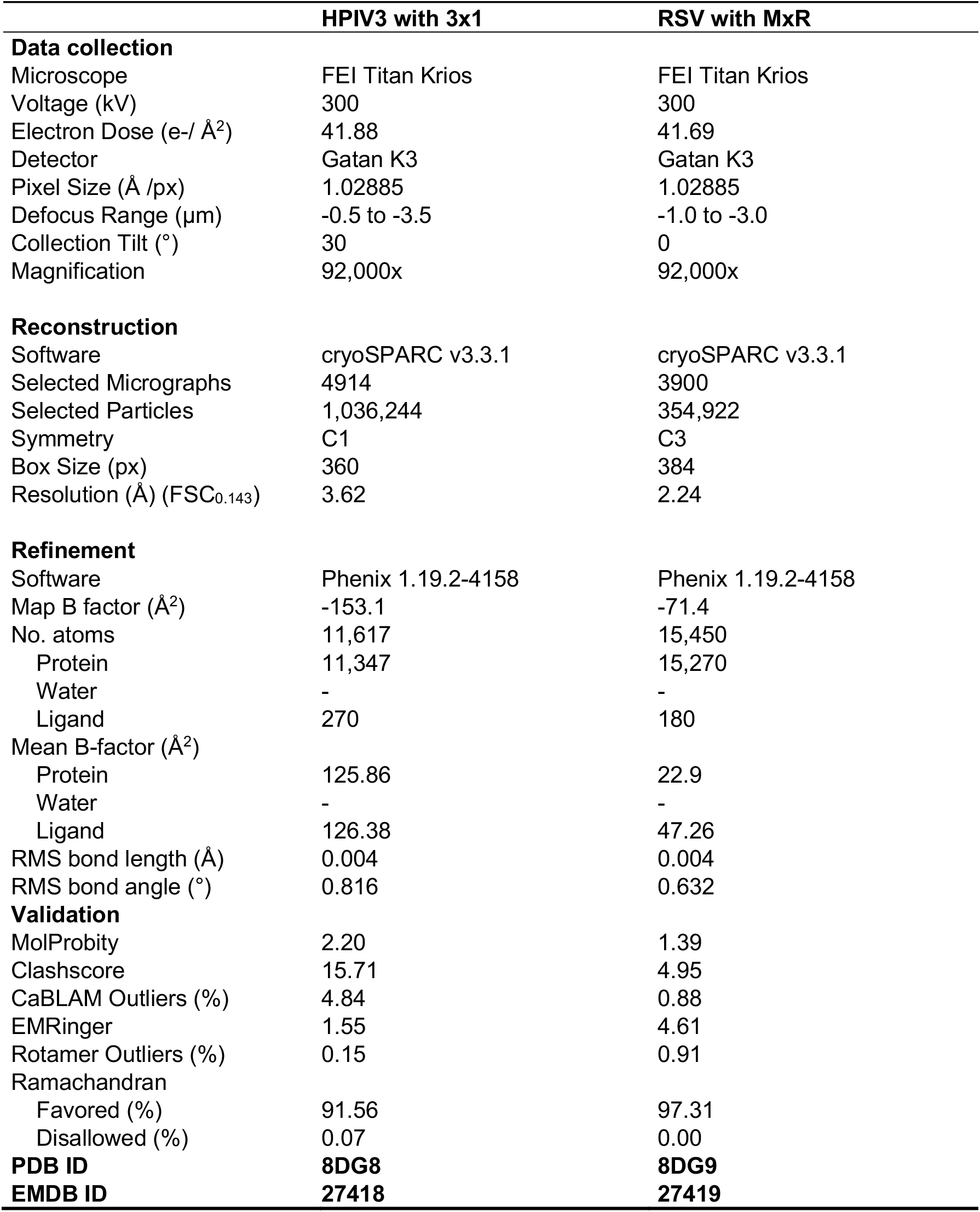
3×1:HPIV3 and MxR:RSV cryo-EM structure statistics

## REFERENCES

1. Giraud-Gatineau A, et al. Comparison of mortality associated with respiratory viral infections between December 2019 and March 2020 with that of the previous year in Southeastern France. Int J Infect Dis 96, 154–156 (2020).

2. Ruckwardt TJ, Morabito KM, Graham BS. Immunological Lessons from Respiratory Syncytial Virus Vaccine Development. Immunity 51, 429–442 (2019).

3. Drysdale SB, Barr RS, Rollier CS, Green CA, Pollard AJ, Sande CJ. Priorities for developing respiratory syncytial virus vaccines in different target populations. Sci Transl Med 12, (2020).

4. Kennedy LB, Li Z, Savani BN, Ljungman P. Measuring Immune Response to Commonly Used Vaccinations in Adult Recipients of Allogeneic Hematopoietic Cell Transplantation. Biol Blood Marrow Transplant 23, 1614–1621 (2017).

5. Ambati A, et al. Evaluation of pretransplant influenza vaccination in hematopoietic SCT: a randomized prospective study. Bone Marrow Transplant 50, 858–864 (2015).

6. Harris AE, Styczynski J, Bodge M, Mohty M, Savani BN, Ljungman P. Pretransplant vaccinations in allogeneic stem cell transplantation donors and recipients: an often-missed opportunity for immunoprotection? Bone Marrow Transplant 50, 899–903 (2015).

7. Boeckh M. The challenge of respiratory virus infections in hematopoietic cell transplant recipients. Br J Haematol 143, 455–467 (2008).

8. Erard V, et al. Airflow decline after myeloablative allogeneic hematopoietic cell transplantation: the role of community respiratory viruses. J Infect Dis 193, 1619–1625 (2006).

9. Agha R, Avner JR. Delayed Seasonal RSV Surge Observed During the COVID-19 Pandemic. Pediatrics 148, (2021).

10. Foley DA, et al. The Interseasonal Resurgence of Respiratory Syncytial Virus in Australian Children Following the Reduction of Coronavirus Disease 2019-Related Public Health Measures. Clin Infect Dis, (2021).

11. Sieling WD, Goldman CR, Oberhardt M, Phillips M, Finelli L, Saiman L. Comparative incidence and burden of respiratory viruses associated with hospitalization in adults in New York City. Influenza Other Respir Viruses 15, 670–677 (2021).

12. Lee N, et al. Burden of noninfluenza respiratory viral infections in adults admitted to hospital: analysis of a multiyear Canadian surveillance cohort from 2 centres. CMAJ 193, E439–E446 (2021).

13. Baker RE, Park SW, Yang W, Vecchi GA, Metcalf CJE, Grenfell BT. The impact of COVID-19 nonpharmaceutical interventions on the future dynamics of endemic infections. Proc Natl Acad Sci U S A 117, 30547–30553 (2020).

14. Corti D, et al. Cross-neutralization of four paramyxoviruses by a human monoclonal antibody. Nature 501, 439–443 (2013).

15. Schiffer JT, Kirby K, Sandmaier B, Storb R, Corey L, Boeckh M. Timing and severity of community acquired respiratory virus infections after myeloablative versus non-myeloablative hematopoietic stem cell transplantation. Haematologica 94, 1101–1108 (2009).

16. Crotty MP, et al. Epidemiology, Co-Infections, and Outcomes of Viral Pneumonia in Adults: An Observational Cohort Study. Medicine (Baltimore) 94, e2332 (2015).

17. Saez-Llorens X, et al. Safety and pharmacokinetics of palivizumab therapy in children hospitalized with respiratory syncytial virus infection. Pediatr Infect Dis J 23, 707–712 (2004).

18. Meissner HC. Viral Bronchiolitis in Children. N Engl J Med 374, 62–72 (2016).

19. Polack FP, Stein RT, Custovic A. The Syndrome We Agreed to Call Bronchiolitis. J Infect Dis 220, 184–186 (2019).

20. White JM, Delos SE, Brecher M, Schornberg K. Structures and mechanisms of viral membrane fusion proteins: multiple variations on a common theme. Crit Rev Biochem Mol Biol 43, 189–219 (2008).

21. Connolly SA, Leser GP, Yin HS, Jardetzky TS, Lamb RA. Refolding of a paramyxovirus F protein from prefusion to postfusion conformations observed by liposome binding and electron microscopy. Proc Natl Acad Sci U S A 103, 17903–17908 (2006).

22. Ngwuta JO, et al. Prefusion F-specific antibodies determine the magnitude of RSV neutralizing activity in human sera. Sci Transl Med 7, 309ra162 (2015).

23. Gilman MS, et al. Rapid profiling of RSV antibody repertoires from the memory B cells of naturally infected adult donors. Sci Immunol 1, (2016).

24. Yin HS, Paterson RG, Wen X, Lamb RA, Jardetzky TS. Structure of the uncleaved ectodomain of the paramyxovirus (hPIV3) fusion protein. Proc Natl Acad Sci U S A 102, 9288–9293 (2005).

25. Moscona A. Interaction of human parainfluenza virus type 3 with the host cell surface. Pediatr Infect Dis J 16, 917–924 (1997).

26. Henrickson KJ. Parainfluenza viruses. Clin Microbiol Rev 16, 242–264 (2003).

27. Boonyaratanakornkit J, et al. Protective antibodies against human parainfluenza virus type 3 infection. MAbs 13, 1912884 (2021).

28. Stewart-Jones GBE, et al. Structure-based design of a quadrivalent fusion glycoprotein vaccine for human parainfluenza virus types 1-4. Proc Natl Acad Sci U S A 115, 12265–12270 (2018).

29. Wen X, Mousa JJ, Bates JT, Lamb RA, Crowe JE, Jr., Jardetzky TS. Structural basis for antibody cross-neutralization of respiratory syncytial virus and human metapneumovirus. Nat Microbiol 2, 16272 (2017).

30. Tang A, et al. A potent broadly neutralizing human RSV antibody targets conserved site IV of the fusion glycoprotein. Nat Commun 10, 4153 (2019).

31. Goodwin E, et al. Infants Infected with Respiratory Syncytial Virus Generate Potent Neutralizing Antibodies that Lack Somatic Hypermutation. Immunity 48, 339–349 e335 (2018).

32. Ottolini MG, Porter DD, Hemming VG, Hensen SA, Sami IR, Prince GA. Semi-permissive replication and functional aspects of the immune response in a cotton rat model of human parainfluenza virus type 3 infection. J Gen Virol 77 (Pt 8), 1739–1743 (1996).

33. Skiadopoulos MH, Surman SR, Riggs JM, Orvell C, Collins PL, Murphy BR. Evaluation of the replication and immunogenicity of recombinant human parainfluenza virus type 3 vectors expressing up to three foreign glycoproteins. Virology 297, 136–152 (2002).

34. Newman JT, et al. Generation of recombinant human parainfluenza virus type 1 vaccine candidates by importation of temperature-sensitive and attenuating mutations from heterologous paramyxoviruses. J Virol 78, 2017–2028 (2004).

35. Liu X, et al. Human parainfluenza virus type 3 expressing the respiratory syncytial virus pre-fusion F protein modified for virion packaging yields protective intranasal vaccine candidates. PLoS One 15, e0228572 (2020).

36. Herfst S, et al. Generation of temperature-sensitive human metapneumovirus strains that provide protective immunity in hamsters. J Gen Virol 89, 1553–1562 (2008).

37. Wen SC, Williams JV. New Approaches for Immunization and Therapy against Human Metapneumovirus. Clin Vaccine Immunol 22, 858–866 (2015).

38. Crowe JE, Jr., et al. Live subgroup B respiratory syncytial virus vaccines that are attenuated, genetically stable, and immunogenic in rodents and nonhuman primates. J Infect Dis 173, 829–839 (1996).

39. Taylor G. Animal models of respiratory syncytial virus infection. Vaccine 35, 469–480 (2017).

40. Schickli JH, Kaur J, Macphail M, Guzzetta JM, Spaete RR, Tang RS. Deletion of human metapneumovirus M2-2 increases mutation frequency and attenuates growth in hamsters. Virol J 5, 69 (2008).

41. Buchholz UJ, et al. Deletion of M2 gene open reading frames 1 and 2 of human metapneumovirus: effects on RNA synthesis, attenuation, and immunogenicity. J Virol 79, 6588–6597 (2005).

42. MacPhail M, et al. Identification of small-animal and primate models for evaluation of vaccine candidates for human metapneumovirus (hMPV) and implications for hMPV vaccine design. J Gen Virol 85, 1655–1663 (2004).

43. Murphy BR, Collins PL. Live-attenuated virus vaccines for respiratory syncytial and parainfluenza viruses: applications of reverse genetics. J Clin Invest 110, 21–27 (2002).

44. Boukhvalova MS, Yim KC, Blanco J. Cotton rat model for testing vaccines and antivirals against respiratory syncytial virus. Antivir Chem Chemother 26, 2040206618770518 (2018).

45. Boukhvalova M, Blanco JC, Falsey AR, Mond J. Treatment with novel RSV Ig RI-002 controls viral replication and reduces pulmonary damage in immunocompromised Sigmodon hispidus. Bone Marrow Transplant 51, 119–126 (2016).

46. Boukhvalova MS, Prince GA, Blanco JC. The cotton rat model of respiratory viral infections. Biologicals 37, 152–159 (2009).

47. Whaley RE, et al. Generation of a cost-effective cell line for support of high-throughput isolation of primary human B cells and monoclonal neutralizing antibodies. J Immunol Methods, 112901 (2020).

48. McLellan JS. Neutralizing epitopes on the respiratory syncytial virus fusion glycoprotein. Curr Opin Virol 11, 70–75 (2015).

49. Corti D, et al. Cross-neutralization of four paramyxoviruses by a human monoclonal antibody. Nature 501, 439–443 (2013).

50. Thomas NJ, Hollenbeak CS, Ceneviva GD, Geskey JM, Young MJ. Palivizumab prophylaxis to prevent respiratory syncytial virus mortality after pediatric bone marrow transplantation: a decision analysis model. J Pediatr Hematol Oncol 29, 227–232 (2007).

51. Hammitt LL, et al. Nirsevimab for Prevention of RSV in Healthy Late-Preterm and Term Infants. N Engl J Med 386, 837–846 (2022).

52. Joyce MG, et al. Crystal Structure and Immunogenicity of the DS-Cav1-Stabilized Fusion Glycoprotein From Respiratory Syncytial Virus Subtype B. Pathog Immun 4, 294–323 (2019).

53. Griffin MP, et al. Single-Dose Nirsevimab for Prevention of RSV in Preterm Infants. N Engl J Med 383, 415–425 (2020).

54. Tixagevimab and Cilgavimab (Evusheld) for Pre-Exposure Prophylaxis of COVID-19. JAMA 327, 384–385 (2022).

55. Biacchesi S, Skiadopoulos MH, Tran KC, Murphy BR, Collins PL, Buchholz UJ. Recovery of human metapneumovirus from cDNA: optimization of growth in vitro and expression of additional genes. Virology 321, 247–259 (2004).

56. Bartlett EJ, et al. Human parainfluenza virus type I (HPIV1) vaccine candidates designed by reverse genetics are attenuated and efficacious in African green monkeys. Vaccine 23, 4631–4646 (2005).

57. Munir S, Le Nouen C, Luongo C, Buchholz UJ, Collins PL, Bukreyev A. Nonstructural proteins 1 and 2 of respiratory syncytial virus suppress maturation of human dendritic cells. J Virol 82, 8780–8796 (2008).

58. McLellan JS, et al. Structure-based design of a fusion glycoprotein vaccine for respiratory syncytial virus. Science 342, 592–598 (2013).

59. Wen X, et al. Structure of the human metapneumovirus fusion protein with neutralizing antibody identifies a pneumovirus antigenic site. Nat Struct Mol Biol 19, 461–463 (2012).

60. Wu X, et al. Rational design of envelope identifies broadly neutralizing human monoclonal antibodies to HIV-1. Science 329, 856–861 (2010).

61. Huang J, et al. Isolation of human monoclonal antibodies from peripheral blood B cells. Nat Protoc 8, 1907–1915 (2013).

62. McGuire AT, et al. Specifically modified Env immunogens activate B-cell precursors of broadly neutralizing HIV-1 antibodies in transgenic mice. Nat Commun 7, 10618 (2016).

63. Johnson RA, Prince GA, Suffin SC, Horswood RL, Chanock RM. Respiratory syncytial virus infection in cyclophosphamide-treated cotton rats. Infect Immun 37, 369–373 (1982).

64. Wu H, et al. Development of motavizumab, an ultra-potent antibody for the prevention of respiratory syncytial virus infection in the upper and lower respiratory tract. J Mol Biol 368, 652–665 (2007).

